# Identification and Expression Analysis of Heat Shock Proteins in Wheat Infected with Powdery Mildew and Stripe Rust

**DOI:** 10.1101/2020.03.26.010801

**Authors:** Hong Zhang, Huan Guo, Guanghao Wang, Changyou Wang, Yajuan Wang, Xinlun Liu, Wanquan Ji

**Author notes:** Corresponding author (H.Z.).

## Abstract

Heat shock proteins (HSPs), which are encoded by conserved gene families in plants, are crucial for development and responses to diverse stresses. However, the wheat (*Triticum aestivum*) HSPs have not been systematically classified, especially those involved in protecting plants from disease. Here, we classified 119 DnaJ (Hsp40) proteins (TaDnaJs; encoded by 313 genes) and 41 Hsp70 proteins (TaHsp70s; encoded by 95 genes) into six and four groups, respectively, via a phylogenetic analysis. An examination of protein structures and a multiple sequence alignment revealed diversity in the TaDnaJ structural organization, but a highly conserved J-domain, which was usually characterized by an HPD motif followed by DRD or DED motifs. The expression profiles of these HSP-encoding homologous genes varied in response to *Blumeria graminis* f. sp. *tritici* and *Puccinia striiformis* f. sp. *tritici*. A quantitative real-time PCR analysis indicated a lack of similarity in the expression of *DnaJ70b, Hsp70-30b*, and *Hsp90-4b* in wheat infected by *B. graminis* f. sp. *tritici*, although the expression levels of these genes were abnormal in the infected resistant and susceptible lines. Furthermore, a direct interaction between DnaJ70 and TaHsp70-30 was not detected in a yeast two-hybrid assay. This study revealed the structure and expression profiles of the HSP-encoding genes in wheat. The resulting data may be useful for future functional analyses and for further elucidating the roles of wheat HSPs during responses to fungal infections.

## Introduction

In the cytoplasm and nucleus, the heat shock response mediates stress-induced transcriptional changes via the increased production of essential protective factors [1] called heat shock proteins (HSPs; also known as molecular chaperones) [2]. On the basis of their molecular weight, HSPs have been classified into several major families, namely Hsp90, Hsp70 (also named DnaK in *Escherichia coli*), Hsp40 (also referred to as DnaJ), Hsp60, and the small Hsps. From yeast to humans, Hsp40s and Hsp70s form chaperone partnerships that are key components of cellular chaperone networks involved in facilitating the correct folding of diverse proteins. The DnaJ molecular chaperones, which represent the crucial partners, bind to and transfer substrate proteins to the Hsp70s to regulate their ATPase activity [3, 4]. The DnaJ proteins comprise the following four domains: N-terminal J-domain, G/F-domain (Gly/Phe-rich region), Zn-binding domain characterized by cysteine repeats, and the C-terminal dimerization domain [5, 6]. Depending on the presence of the G/F-rich region and/or the cysteine repeats, a DnaJ/Hsp40 protein is categorized as type I, II, or III [7]. The Hsp70/Hsp40/NEF (nucleotide exchange factor) system assists in intracellular protein refolding and helps maintain proteostasis in healthy and stressed cells [7-10]. Additionally, Hsp70 functions cooperatively with the highly conserved Hsp90 via co-chaperones [11, 12], resulting in the assembly, maturation, stabilization, and activation of key signalling proteins, including protein kinases and transcription factors in eukaryotic cells [13, 14]. These observations suggest there is an important cycle of functions related to the interaction between Hsp70 and the J-domain proteins Hsp40 and Hsp90 [15, 16]. The Hsp90 molecular chaperone also ensures proteins are maintained in their active conformations [17].

Plants have no capacity to escape from adverse environments, and their growth and production are severely affected by abiotic and biotic stresses. To ensure they can successfully propagate, plants have developed multiple defence mechanisms that allow them to detect pathogens and induce rapid responses through innate immunity surveillance systems [18, 19]. Genes encoding HSPs are reportedly differentially expressed in various plant species exposed to abiotic and biotic stresses. During plant-pathogen interaction, the *AtHsp90.1* gene is required for the full *RESISTANT TO P. SYRINGAE 2* (*RPS2*)-mediated resistance against *Pseudomonas syringae* pv. *tomato* DC3000 (*avrRpt2*), and its function is closely associated with *RAR1* (required for Ml<u>a</u>12 resistance) and *SGT1* (<u>s</u>uppressor of the <u>G</u>2 allele of skp1) via chaperone activities [20]. Similarly, suppressing *TaHsp90.2* or *Hsp90.3* expression in wheat decreases the hyper-sensitive resistance to the stripe rust fungus [21]. Previous studies also confirmed that Hsp70s and Hsp40s are involved in various plant disease resistance, such as NtMPIP1 [22], GmHSP40.1 [23], OsDjA6 [24]. The importance of the proteins encoded by these genes in responses to various environmental stimuli [25-27] and their dynamic interplay with the chaperone machinery suggest that targeting Hsp90 and its respective co-chaperones may be an effective method for characterizing the mechanisms underlying the resistance to diverse plant diseases.

We previously constructed two protein–protein interaction networks based on a weighted gene co-expression network analysis (WGCNA) to clarify the mechanism mediating wheat responses to the pathogens causing stripe rust (*Puccinia striiformis* f. sp. *tritici*; *Pst*) and powdery mildew (*Blumeria graminis* f. sp. *tritici*; *Bgt*) [28]. Both of these networks predicted that Hsp70 proteins represent a key hub node because they interact with some splicing regulators, transcription factors, and resistance (*R*) genes, including the disease resistance-related *RPP13, RPS2* analogues, the pathogenesis-related protein 1 gene (*PR1*), and a non-host resistance gene (*NHO1*) [28]. In this study, we identified HSPs and their genes in wheat (*Triticum aestivum*). Moreover, the expression of these genes following powdery mildew and stripe rust infections was analysed with RNA-sequencing (RNA-seq) and quantitative real-time polymerase chain reaction (qRT-PCR) assays. Furthermore, the interactions among selected HSPs were assessed in a yeast two-hybrid (Y2H) assay.

## Materials and Methods

### Plant materials and pathogen stress treatment

The N9134 winter wheat cultivar, which is resistant to all *Bgt* races in China, was crossed seven times with a susceptible recurrent parent, Shaanyou225 (SY225), to obtain a resistant line with the SY225 background. This new resistant line was named SY225/7 × PmAS846. To eliminate the influence of inhomogeneity, a BC_7_F_1_ resistant plant was self-crossed to generate the contrasting BC_7_F_2_ homozygous lines, which differ only regarding *PmAS846* on chromosome 5BL. Additionally, *Bgt* isolate E09 was maintained on SY225 wheat plants. The SY225/7 × PmAS846 and SY225 plants were cultivated in soil in a growth chamber at 18 °C with a 16-h light/8-h dark photoperiod. Seedlings at the 3-leaf stage were inoculated with *Bgt* conidia from SY225 seedlings that were inoculated 10 days earlier. The inoculated N9134 leaves were harvested at 6, 12, 24, 36, 48, 72, and 96 hours post-inoculation (hpi), after which they were immediately frozen in liquid nitrogen and stored at −80 °C. Subsequent analyses were completed with three biological replicates. For the genome-wide transcription analysis, 7-day-old seedlings were inoculated with *Bgt* E09 or *Pst* race CYR 31 conidia. The inoculated N9134 leaves were harvested at 0, 24, 48, and 72 hpi as previously described [29]. The non-inoculated sample (0 hpi) was used as the control.

### Identification and sequence analyses of wheat heat shock proteins

In order to obtain detailed information regarding wheat HSPs, we downloaded all available sequences for proteins annotated with ‘Hsp’ in the EnsemblPlants database (http://plants.ensembl.org/Triticum_aestivum/Info/Index). After screening the NCBI database with the BLASTP (Protein Basic Local Alignment Search Tool) algorithm for sequence matches (80% identity as the threshold), the HSP proteins were sorted and classified. Redundant sequences were manually removed and the remaining sequences were analysed with the NCBI Conserved Domain database (http://www.ncbi.nlm.nih.gov/Structure/cdd/wrpsb.cgi) and the Pfam database to detect conserved protein domains and identify candidate *HSP* genes. The motifs of the encoded HSP proteins were detected using the motif-based sequence analysis program (MEME) [30], whereas a phylogenetic neighbour-joining tree was constructed for the HSP proteins with the molecular evolutionary genetics analysis (MEGA) program (version 6.0) [31].

### Quantitative real-time PCR analysis

The differentially expressed HSP-encoding genes that involved in wheat responses to *Bgt* and *Pst* infections we screened using the RNA-seq data (PRJNA243835) from the fungus-inoculated N9134 (resistant to *Bgt*-E09 with *PmAS846* and CYR31 with *YrN9134*). The detailed RNA-seq protocol was described as the previously report [28]. The reads per kilobase of exon model per million of aligned readings (RPKM) values were used to examine the gene expression level distribution for each transcription in sample. In addition, the expression profiles of three spcific HSP-encoding genes in the infected wheat leaves of the contrasting NILs were analysed by a SYBR Green-based qRT-PCR with cDNA prepared from samples collected at 6, 12, 24, 36, 48, 72, and 96 hpi. The uninoculated plant samples at the same time points were set as the controls. Three independent biological replicates were prepared for each time point. The qRT-PCR was completed with the FastKing RT Kit (with gDNase) (TIANGEN, Beijing) and the QuantStudio™ 7 Flex Real-Time PCR System (Life Technologies Corporation, USA). Primers specific for the examined HSP-encoding genes (**Table S1**) and *TaActin* (i.e., internal reference) were designed with the Primer 5 program, and were used to analyse gene expression levels. The 20-µL reaction volume comprised 10 µL 2× SYBR Green PCR Master Mix (Takara, Dalian, China), 0.2 µM each primer, and 2 µL template (6× diluted cDNA from leaf samples). The PCR program was as follows: 95 °C for 10 s; 40 cycles of 95 °C for 5 s and 60 °C for 31 s. For each sample, reactions were completed in triplicate, with three non-template negative controls. Amplified products were analysed with melting curves, which were generated at the end of the amplification. A standard 2^−ΔΔCT^ method was used to quantify relative gene expression levels.

### Transcriptional activity analysis and yeast two-hybrid assays

The expression of *HIS3, ADE2*, and *LacZ* reporter genes was examined with yeast AH109 transformants on SD/−Trp/−His and SD/−Trp/−Ade media (Clontech Inc., USA). The fix composition of SD plates contained 0.17 g yeast nitrogen base (YNB), 0.5 g ammonium sulfate, 2 g glucose, 2 g agar and 1 ml 10×Dropout (-His –Leu –Trp –Ura) in 100 ml volume. The full-length open reading frame of *TaHsf-B1b* as well as the N-terminal domain and the C-terminal regulatory domain were amplified by PCR with specific primers containing a homologous recombination arm (**Table S1**). To easily evaluate transcriptional activity, the transformants were suspended in sterile distilled water, after which 10-fold serial dilutions were prepared. Finally, 3-μL aliquots of each dilution were used to inoculate SD/−Trp/−His/−Ade medium and SD/−Trp/−His medium with X-α-Gal (Clontech). The inoculated media were incubated for 4 days at 30 °C. The MatchMaker yeast two-hybrid system (Takara, Dalian, China) was used to evaluate the interactions among TaHsp70-30b, TaHsp90-4b, and TaDnaJ70b. The *TaHsp70-30b, TaHsp90-4b*, and *TaDnaJ70b* coding sequences were subcloned into the pGBKT7 (DNA-binding domain, BD) and pGADT7 (activation domain, AD) vectors. Additionally, the pGADT7-Sfi I three-frame primary cDNA library was constructed by Takara (Dalian, China) using leaf samples from N9134 after infected with stripe rust fungi at 24 and 48 hours. The AD library was screened for protein interactions by mating pGBKT7:TaHsp70-30b bait plasmid, while specific AD and BD recombinant plasmid pairs were used to co-transform yeast strain AH109 cells according to the Yeast Protocols Handbook (Takara, Dalian, China).

### Statistical analysis

Mean values and standard errors were calculated with Microsoft Excel software. Student’s *t*-test was completed with the SPSS 16.0 program to assess the significance of any differences between the control and treated samples or between time-points. The threshold for significance was set at *P* < 0.01.

## Results

### Identification of wheat heat shock protein 40 (DnaJ) family proteins

In order to elucidate the mechanism mediating wheat resistance to *Pst* and *Bgt*, we recently predicted the key genes based on a WGCNA and a transcriptome–proteome associated analysis. The predicted genes included TraesCS5B02G374900, which had the most significant connectivity to other genes (value reaching 2,676) [28] and was annotated as encoding a DnaJ-like protein. Screening the *PmAs846* physical map, with the Chinese Spring wheat genome used as a reference sequence, indicated that TraesCS5B02G374900 is located between the co-segregation markers *BJ261635* with *XFCP620* flanking *PmAS846* [32]. To further evaluate the genomic distribution and function of HSPs in wheat defences against pathogens, RNA-seq data were examined for a genome-wide identification of HSP-encoding genes in *Triticum aestivum*, including genes responsive to fungi. The resulting data confirmed the presence of 119 genes encoding proteins with a characteristic J-domain in the wheat genome as well as 13 genes for proteins with cysteine repeats without a J-domain and 12 genes for proteins with a J-domain, but lacking an HPD motif. These three gene groups were designated as *TaDnaJ, TaDnaJ-CR*, and *TaDnaJL*, respectively. These proteins were encoded by 376 wheat genes. Specific details are provided in Supplemental Table S2. Interestingly, the J-domain was usually accompanied by a domain with the <u>D</u>XXX<u>R</u>XXX<u>D</u> motif or a long DED repeat motif (RRRYGLA<u>DED</u>LDRYRXYLNXX<u>DED</u>DWF) (**Figure 1**). The DnaJ C-terminals usually harboured a GK-rich domain characterized by a glycine and lysine interval zipper or a domain with a <u>WA</u>X<u>Y</u> motif.

**Figure 1.**
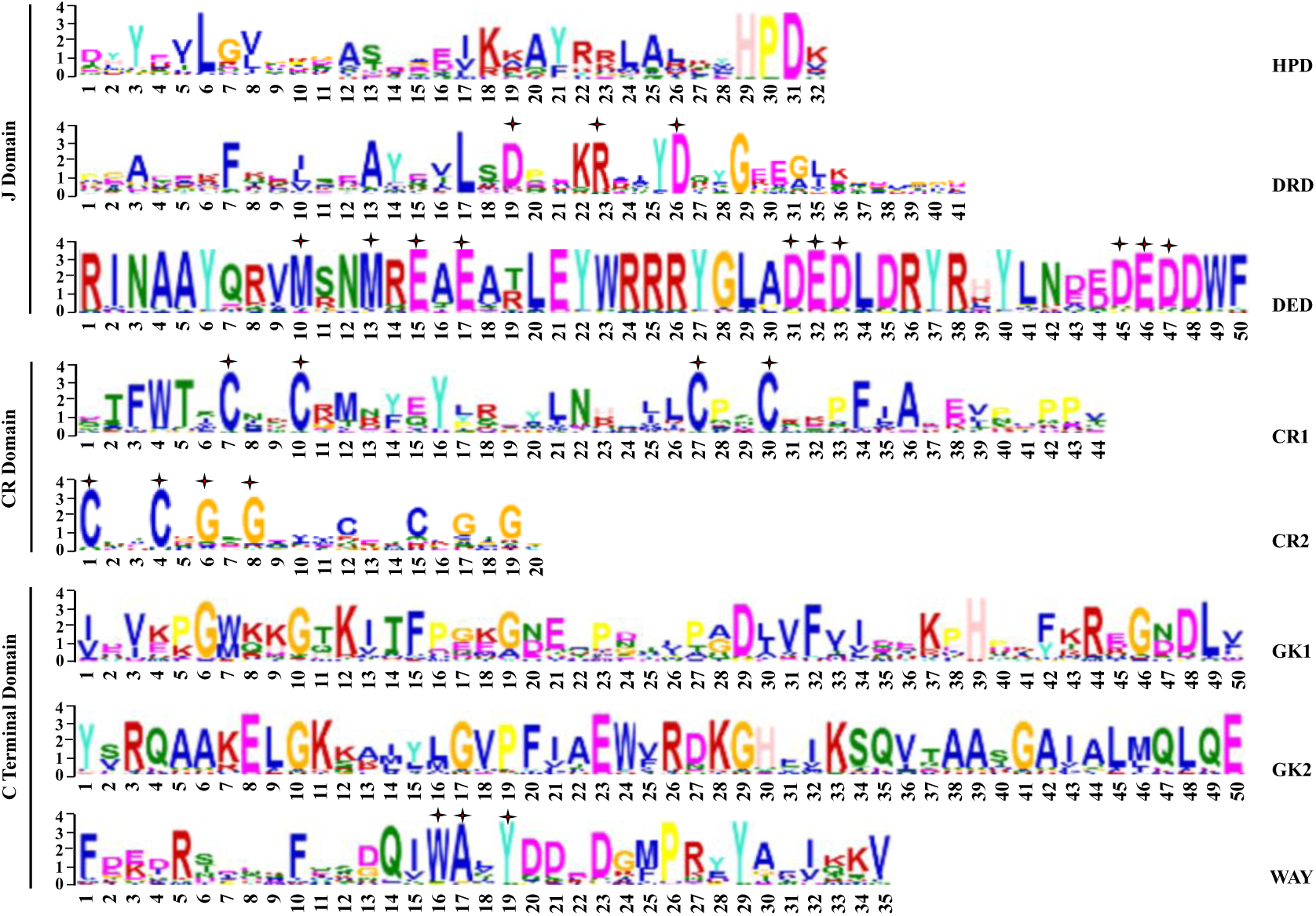
The conserved functional domains and motifs in DnaJ proteins of wheat. The characterization of amino acids were identified by multiple alignment using MEME software. The conserved amino acids of motifs were marked with star signal above the letters. The name of motifs were indicated at the right of each sequence. J domain: KKAYRRLALKY**HPD**; DRD domain: AEEKFQEIQEAYEVLS**D**PEK**R**ALY**D**QYG; DED domain: RINAAYQRV**M**SN**M**R**E**A**E**ATLEYWRRR YGLA**DED**LDRYRHYLNDE**DED**DWF; CR domain 1: **C**PT**C**R**G**S**G**EVV**C**DT**C**N**G**T**G**G; CR domain 2: FWTS**C**NK**C**RMNYZYPREYLNRALL**C**PS**C**RK; GK zipper 1: IDVKP**G**WKK**G**T**K**ITFPGK**G**BEAPD; GK zipper 2: **GK**KAIYL**G**VPFIAEWVRD**KG**HFI**K**SQVTAAS**G**; WAY domain: **F**DED**R**SEEK**F**QSDQI **WA**V**Y**DD EDGMPRY**Y**ARIKKV.

According to the present classification of DnaJ, the N-terminal J-domain of class I DnaJ proteins was followed by a G/F-rich region, four repeats of the CXXCXGXG-type zinc-finger, and a C-terminal extension [33]. However, sequence alignment results indicated that only 17 wheat DnaJ proteins (not including 13 CR domain-containing DnaJ proteins) had a cysteine repeat domain. Of these proteins, nine comprised four repeats of CXXCXGXG and the other eight proteins consisted of two repeats of the CXXC motif. Similarly, only 17 proteins contained a G/F-domain, which was replaced by AP-, SP-, AS-, GS-, or GR-rich domains in the other wheat DnaJ proteins. Additionally, in this study, we detected 119 wheat DnaJ proteins that were encoded by 313 genes (**Supplemental Table S2)**. We observed that Class II proteins present variable domains, whereas most of the Class III proteins consist of domains with characteristic DRD or DED motifs. This means that large structural and potential functional differences exist within and between traditional Classes II and III of DnaJ. Considering the fact that the presence (type II) or absence (type III) of the G/F-domain has led to some ambiguity [33], the 119 wheat DnaJ proteins were further classified in six subfamilies (**Figure 2**) on the basis of the characterized domains. Type I DnaJ proteins were characterized by a DRD motif, whereas type IV and type V DnaJ proteins harboured the DED and WAY motifs, respectively. The type II DnaJ proteins had a classical zinc-finger domain. The J-domain was accompanied by an uncharacterized N-terminal on the type-III DnaJ proteins.

**Figure 2.**
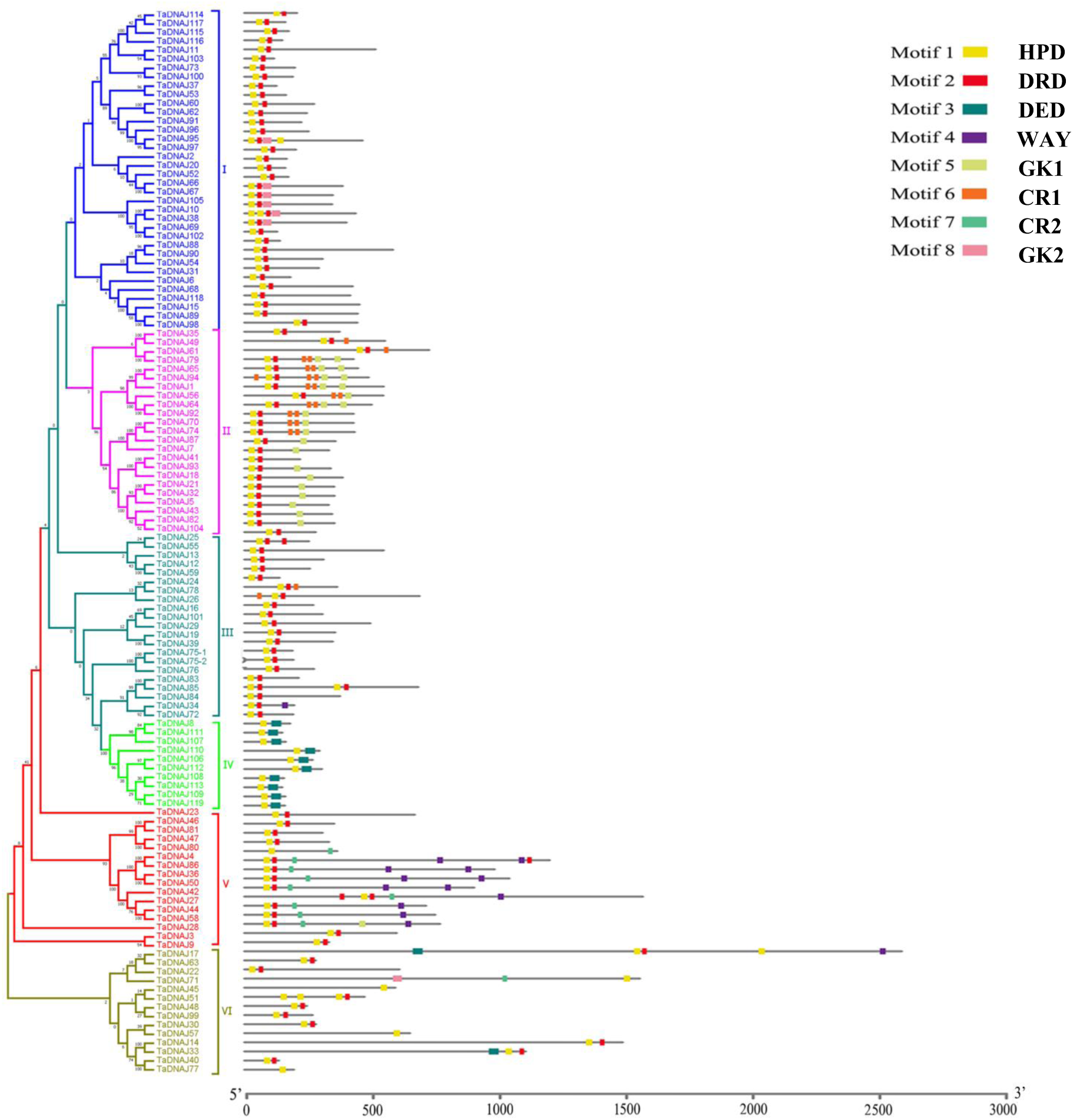
Phylogenetic tree, conserved motifs, and sequence structure of 119 TaDnaJ proteins. The conserved motifs were listed as differential colored box. The sequence structure of TaDnaJ proteins were represented with the mixed thin line and color boxes, while the scale was given in the bottom of panel.

### Identification of the wheat 70- and 90-kD heat shock proteins

We detected 41 Hsp70 proteins (encoded by 95 genes) as well as two proteins containing partial Hsp70 domains, two Hsp70-like proteins, and six Hsp90 proteins (encoded by 18 genes) in the wheat reference genome. Specific details are listed in **Supplemental Tables S3 and S4**. In wheat, the Hsp70 proteins are more conserved than the DnaJ proteins. The six domains detected in 41 unabridged Hsp70 proteins are presented in **Figure 3**. Of these domains, the NX<u>DEAVA</u>, DXX<u>LGG</u>X<u>D</u>, and <u>TPS</u>X<u>VA</u>F motifs were detected in nearly all members. The repetition of the C-terminal <u>E</u>X<u>E</u> motif (wherein X represents glycine, isoleucine, alanine, aspartic acid, leucine, or phenylalanine) was used to classify the Hsp70 proteins in four subfamilies (**Figure 3)**. The type IV Hsp70 proteins lacked the EXE motif, whereas the type II Hsp70 proteins had more EXE repeats than the type I and type III proteins. Interestingly, the type II Hsp70 proteins contained an EEVD pattern at the C-terminal, unlike the type III proteins, which comprised a C-terminal HDEL pattern. Another difference between the type I and type III proteins was the presence of the non-typical linker motif 7 in the type III proteins (**Figure 3**). We detected far fewer Hsp90 proteins than Hsp70 and DnaJ proteins. However, the wheat Hsp90 proteins were observed to contain very similar ED-enriched domains, usually accompanied by a leucine zipper.

**Figure 3.**
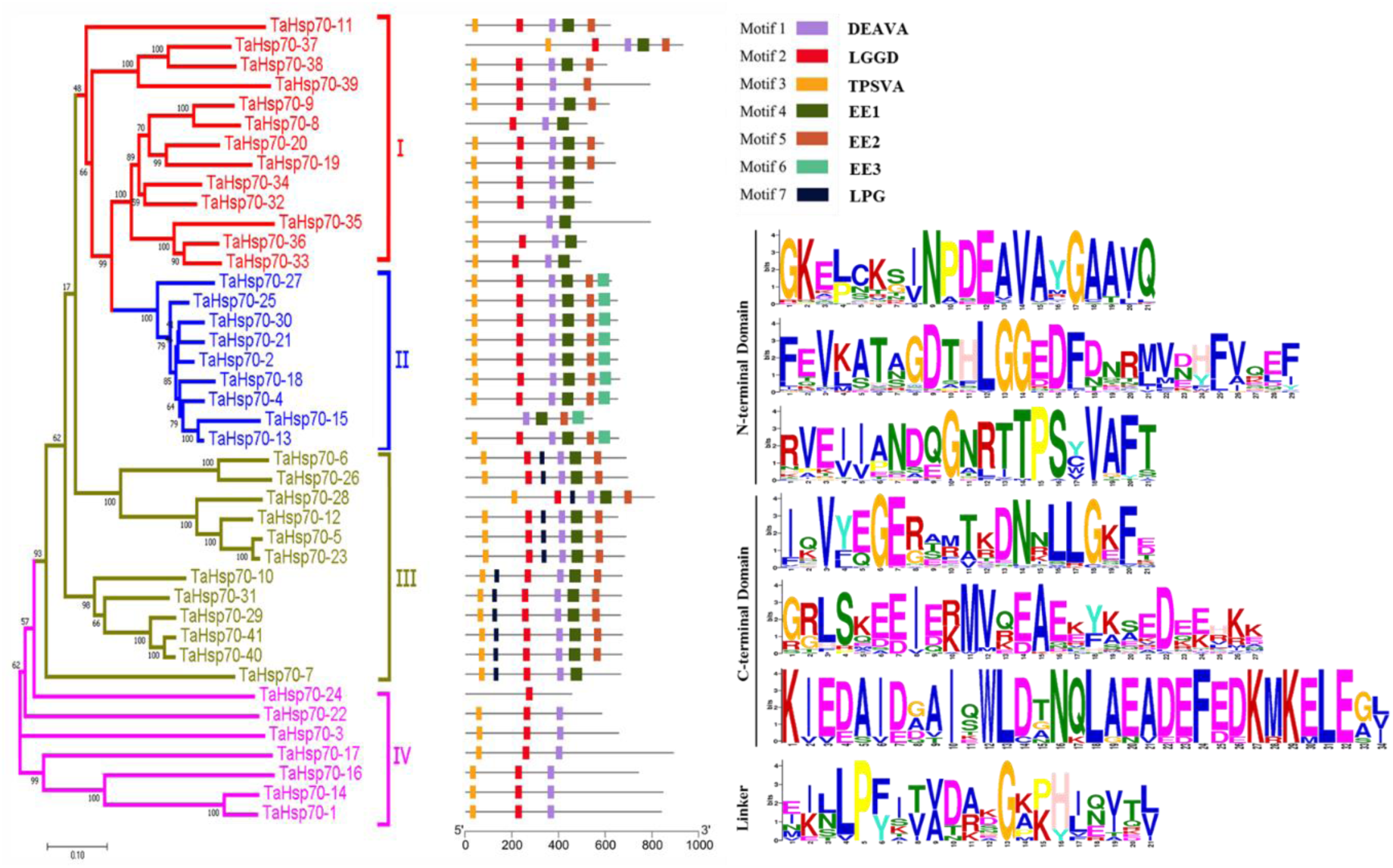
Phylogenetic tree, conserved motifs, and sequence structure of Hsp70 proteins in wheat. The conserved motifs were listed as differential colored box, and their names and sequences were orderly shown in the right. The gene structure of TaHsp70 proteins were represented with the mixed thin line and color boxes, while the scale was given in the left bottom of panel. The characterization of amino acids were identified by multiple alignment using MEME software. Motif 1: GKELCKSI**N**P**DEAVA**YGAAVQ; motif 2: FEVKATAG**D**TH**LGG**E**D**FDNRMVDHFV; motif 3: RVEIIANDQGNRT**TPS**Y**VAF**TDTER; motif 4: IQVY**EGE**RAMTKDNNLLGEFE; motif 5: GRLSKE**EIE**KMVQ**EAE**KYKS**EDE** EHKK; motif 6: KIEDAIDGAIQWLDTNQLAEAD**EFE**DKMK**ELE**GVCNP; motif 7 (seem to be linker): EIL**LP** FITVDAK**G**KPHIQVTL.

### Expression of heat shock protein-coding genes in wheat–Bgt and –Pst interactions

In order to identify the HSP-encoding genes involved in wheat responses to *Bgt* and *Pst* infections, we screened the RNA-seq data (PRJNA243835). In 10-day-old N9134 seedlings inoculated with *Bgt*, 45 *DnaJ* family genes encoding 20 HSPs were identified as differentially expressed genes (relative to the control expression level) (**Figure 4;** Supplemental **Table S2**). Moreover, 14 *Hsp70* genes (encoding seven Hsp70 proteins) and six *Hsp90* genes (encoding three Hsp90 proteins) were identified in the fungus-inoculated N9134 seedling leaves (Supplemental **Table S3 and S4**). Notably, the number of DNAJs was three times than that of Hsp70s, while it was near to seven times than Hsp90s. Conversely, the ratio of the over lapped genes was steeply increased, which reach to 20%, 57.1% and 66.7% for DnaJs, Hsp70s and Hsp90s, respectively. Considering wheat comprises a polyploid genome, we further analysed the differentially expressed genes encoding the same HSP on partially homologous chromosomes. The results indicated that the differentially expressed *Hsp* genes from the partially homologous chromosomes had nearly coincident expression patterns, although the expression levels differed (**Figure 5** and **Supplemental Figure S1**). For example, the expression of *TaDnaJ7-1A, TaDnaJ7-1B*, and *TaDnaJ7-1D* increased at 24 h post-inoculation (hpi) in *Bgt*-infected plants, after which the expression sharply decreased at 48 and 72 hpi (**Supplemental Figure S1**). The transcription levels of *TaDnaJ7-1A* and *TaDnaJ7-1D* were 2.5- and 2.2-fold higher than that of *TaDnaJ7-1B*, respectively. Additionally, *TaDnaJ86-6A, TaDnaJ86-6B*, and *TaDnaJ86-6D* expression levels were not up-regulated until 48 hpi, after which they continued to increase, peaking at 72 hpi. Subsequently, we analysed the level of protein expression from previous quantitative proteomic data (PeptideAtlas: PASS00682 and PASS00999) [28, 34], and found that TaDnaJ7, TaDnaJ70, TaHsp70-4, TaHsp70-25, TaHsp70-30 and TaHsp90-4 proteins were also differentially accumulated with >1.2-fold change in abundance and *t*-test *p*-value < 0.05 (FDR < 1%) in fungi infected leaves comparing to the uninfected control.

**Figure 4.**
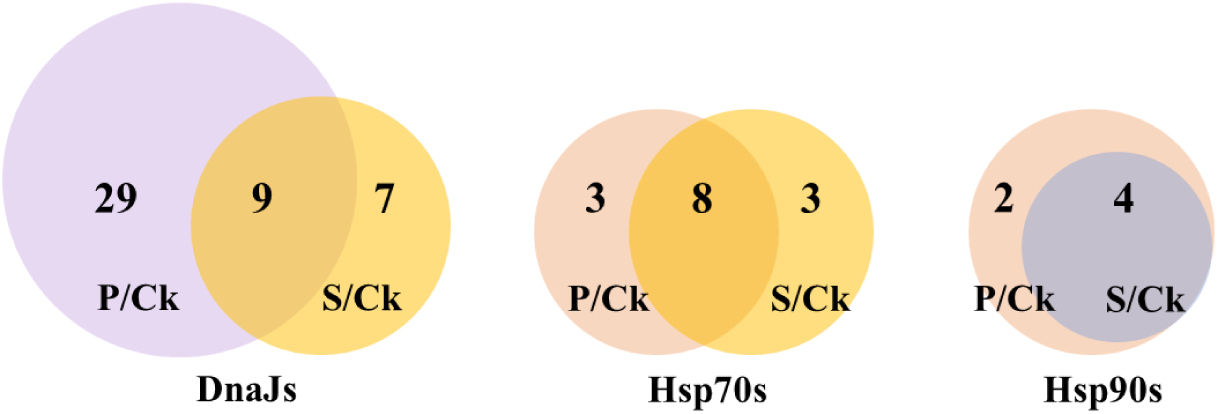
Venn diagram for the overlap differentially expressed Hsp-encoding genes identified with *Bgt* and *Pst* treatments. The number of specific and overlapped genes were given.

**Figure 5.**
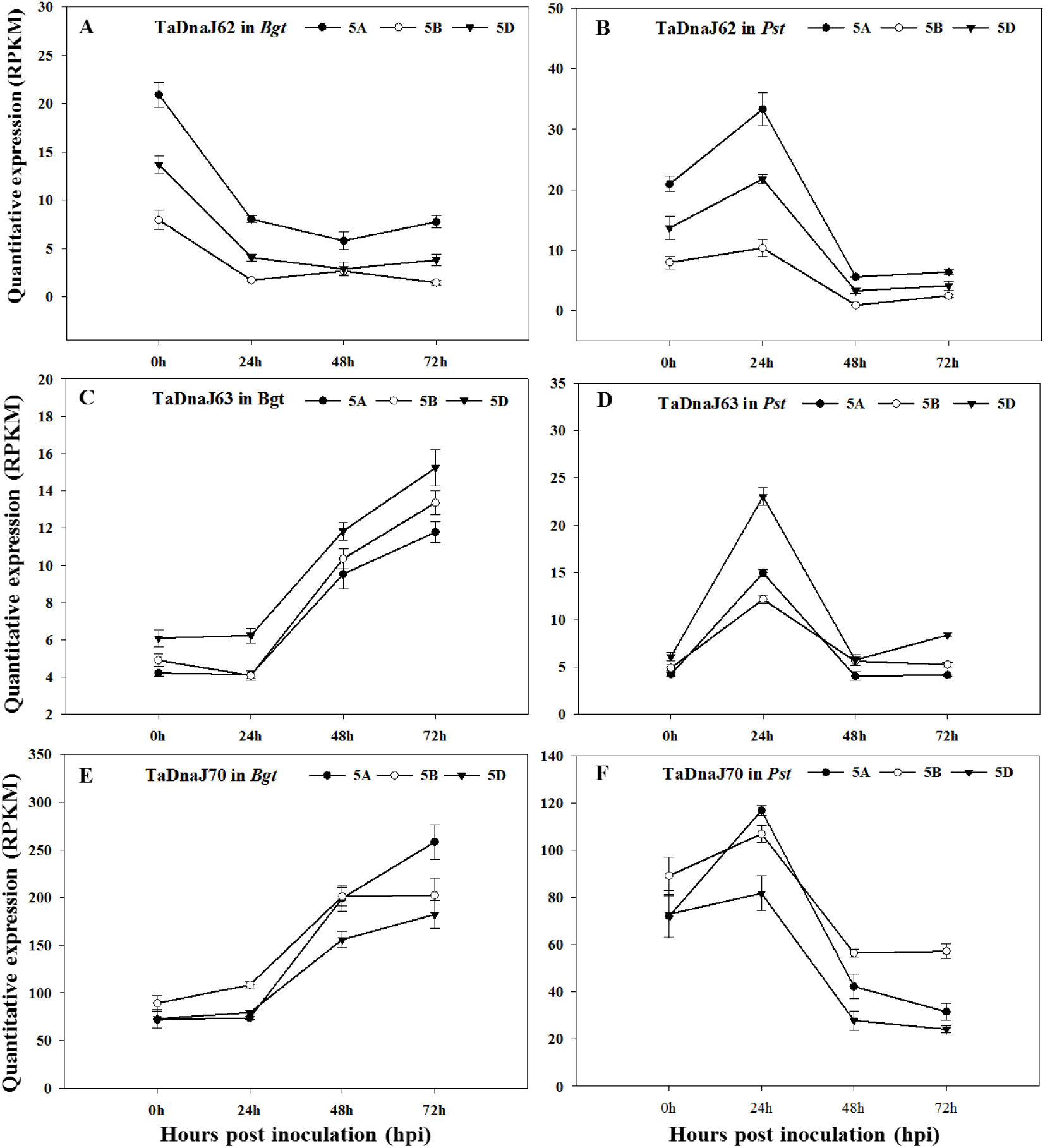
Expression patterns of differentially expressed DnaJ-encoding genes in N9134 infected by stripe rust and powdery mildew pathogen. Gene expression levels were assessed by transcript accumulation analysis. The mean expression value was calculated from three independent replicates. Line charts mean the gene expression of homologue HSP-encoding genes in wheat infected by stripe rust and powdery mildew. The number 0, 24, 48 and 72 indicated the timepoints after infection. A,C and E represent *Bgt* E09 inoculation condition; B, D and F represent *Pst* CYR 31 inoculation. The name of DnaJ-encoding genes and corresponding chromosome were listed in the top of each panel.

The overlapping differentially expressed *Hsp* genes exhibited the complete opposite expression patterns in response to the *Bgt* and *Pst* infections (**Figure 5**). Specifically, the transcription of *TaDnaJ62-5A, TaDnaJ62-5B*, and *TaDnaJ62-5D* were repressed at 24 hpi in the *Bgt*-infected plants, but induced at the same time point in the *Pst*-infected plants. The *TaDnaJ70* expression level was up-regulated at 48 and 72 hpi in the *Bgt*-infected plants, but down-regulated in the *Pst*-infected plants. The *TaHsp70-1, TaHsp70-4, TaHsp70-25*, and *TaHsp90-6* expression levels were induced at 24 hpi in *Bgt*-infected plants, whereas they were repressed at 24 hpi in the *Pst*-infected plants (**Figure 6** and **Supplemental Figure S2**). These observations supported our previous view that distinct genes and regulatory networks are activated in wheat to counter the adverse effects of *Bgt* and *Pst* infections [29, 35].

**Figure 6.**
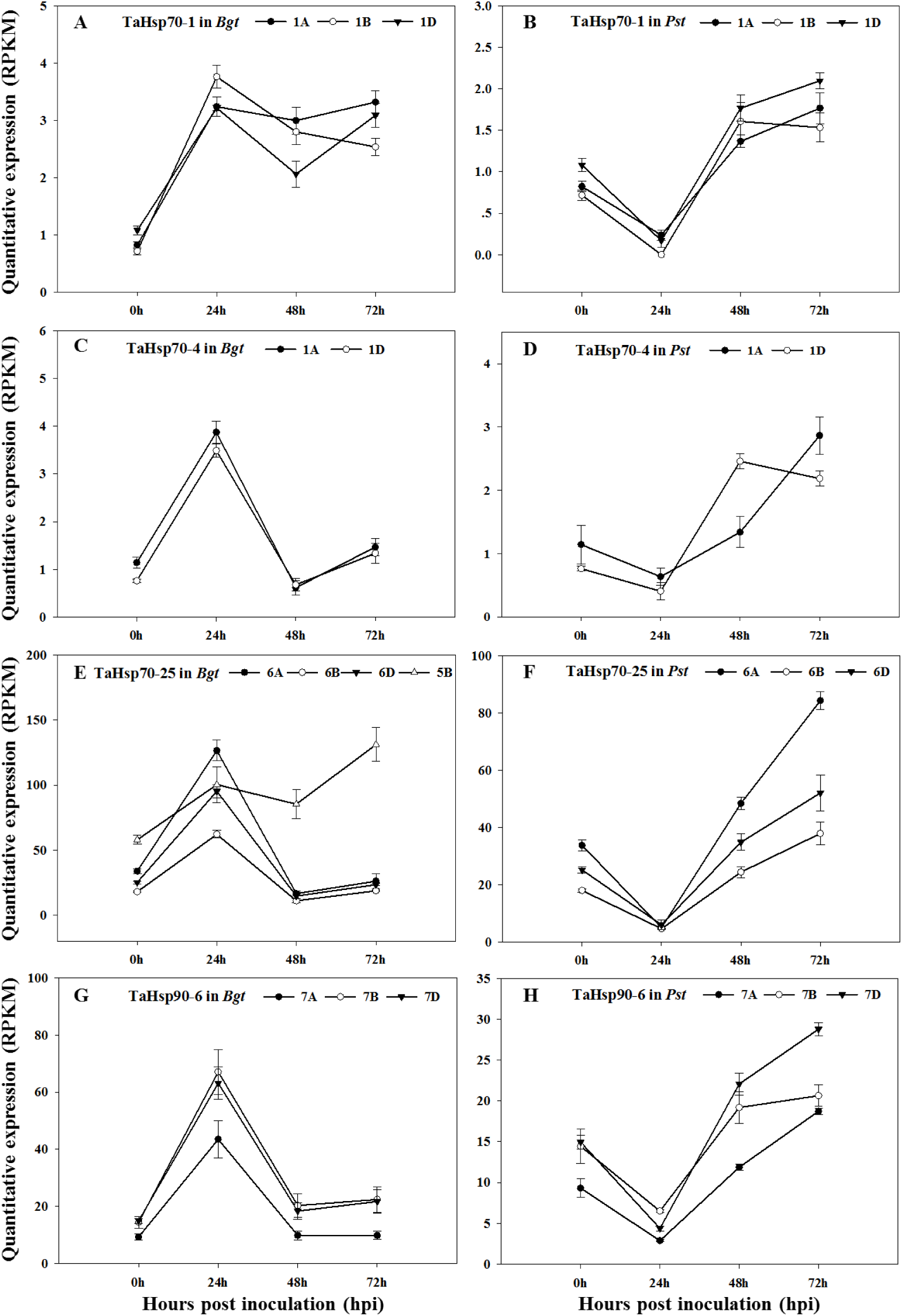
Expression patterns of differentially expressed Hsp70- and Hsp90-encoding genes in N9134 infected by stripe rust and powdery mildew pathogen. Gene expression levels were assessed by transcript accumulation analysis. The mean expression value was calculated from three independent replicates. Line charts mean the gene expression of homologue HSP-encoding genes in wheat infected by stripe rust and powdery mildew. The number 0, 24, 48 and 72 indicated the timepoints after infection. A, C, E and G represent *Bgt* E09 inoculation condition; B, D, F and H represents *Pst* CYR 31 inoculation. The name of HSP-encoding genes and corresponding chromosome were listed in the top of each panel.

In addition to the wheat HSPs, we identified three *Bgt*-*DnaJ* homologues (accession numbers KE373410.1, KE374995.1, and KE375122.1), one *Bgt*-*Hsp60* (CAUH01004772.1), four *Bgt*-*Hsp70* (JQ917466.1, EPQ67599.1, EPQ66911.1, and KE375027.1), and two *Bgt*-*Hsp90* (KE374996.1 and EPQ62547.1) genes (**Table 1**). These fungal HSP-encoding genes were detected after plants were infected (Supplemental **Figure S3**) and had varying expression patterns. The Hsp70-encoding gene JQ917466.1 and Hsp90-encoding gene KE374996.1 were most highly expressed at 48 hpi, whereas the expression of the *DnaJ* homologue KE374995.1 stably increased after plants were infected. These results implied that the HSPs may contribute to the fungal infections of susceptible host plants.

**Table 1.** The detected Bgt-HSP encoding genes in N9134 after powdery mildew infection.

| Encoding protein | Accession No. | Accession No. | Accession No. | Accession No. |
| --- | --- | --- | --- | --- |
| **Bgt-DNAJ** | KE373410.1 | KE374995.1 | KE375122.1 |  |
| **Bgt-Hsp60** | CAUH01004772.1 |  |  |  |
| **Bgt-Hsp70** | JQ917466.1 | EPQ67599.1 | EPQ66911.1 | KE375027.1 |
| **Bgt-Hsp90** | KE374996.1 | EPQ62547.1 |  |  |

### Co-expression of TaHsp70-30b, TaHsp90-4b, and TaDnaJ70b in wheat–Bgt interactions

To assess the expression and function of HSP-encoding genes, *TaDnaJ70b, TaHsp70-30b, TaHsp90-4b*, and a wheat heat shock transcription factor gene, *TaHsf-B1b*, all of which are located on chromosome 5B, were analysed regarding their expression level in the *Bgt*-infected leaves of resistant line N9134R (SY225/7 × PmAS846, a Shaanyou225 backcross line carrying *PmAS846* from N9134) and the contrasting susceptible N9134S NILs. Because of the repetition in the wheat genome sequence, we used four pairs of specific primers targeting the 5′ terminal sequences to target the genes of interest on chromosome 5B. The expression profiles of *TaHsf-B1b, TaDnaJ70b, TaHsp70-30b*, and *TaHsp90-4b* in the inoculated N9134R and N9134S plants are presented in **Figure 7**. Following the inoculation with *Bgt*, the expression levels of all tested genes were dysregulated at 6 hpi significantly, whereas they were oppositely down- and up-regulated in the N9134R and N9134S plants respectively. There was a subsequent steep increase in the expression level at 12 hpi in the N9134R plants. The *TaHsf-B1b, TaDnaJ70b*, and *TaHsp70-30* expression levels at 6 hpi were higher than that in the mock-inoculated control N9134S plants. Afterward, the high expression levels of *TaDnaJ70b, TaHsp70-30b* and *TaHsp90-4b* fluctuated irregularly in N9134S plants, with *TaDnaJ70b* expression peaking at 24 and 96 hpi. Conversely, in N9134R plants, *TaHsf-B1b, TaDnaJ70b*, and *TaHsp70-30b* were relatively stably expressed at low levels, with only slight and consistent fluctuations from 12 to 96 hpi. The highest *TaHsp90-4b* expression levels were observed at 96 hpi in N9134S plants, although they were not significantly higher at other time points than the corresponding expression levels in control (**Figure 7**) in resistance background. Thus, there were no co-expression relationships between *TaHsp90-4b* with *TaDnaJ70b* and *TaHsp70-30b*, while the gene expression pattern of *TaHsf-B1b* was very similar to that of *TaDnaJ70b* and *TaHsp70-30b*. Additionally, the expression levels of the DnaJ and HSP70-encoding genes were generally higher in the susceptible plants than in the resistant plants, but the expression of *HSP90-4b* is just the reverse.

**Figure 7.**
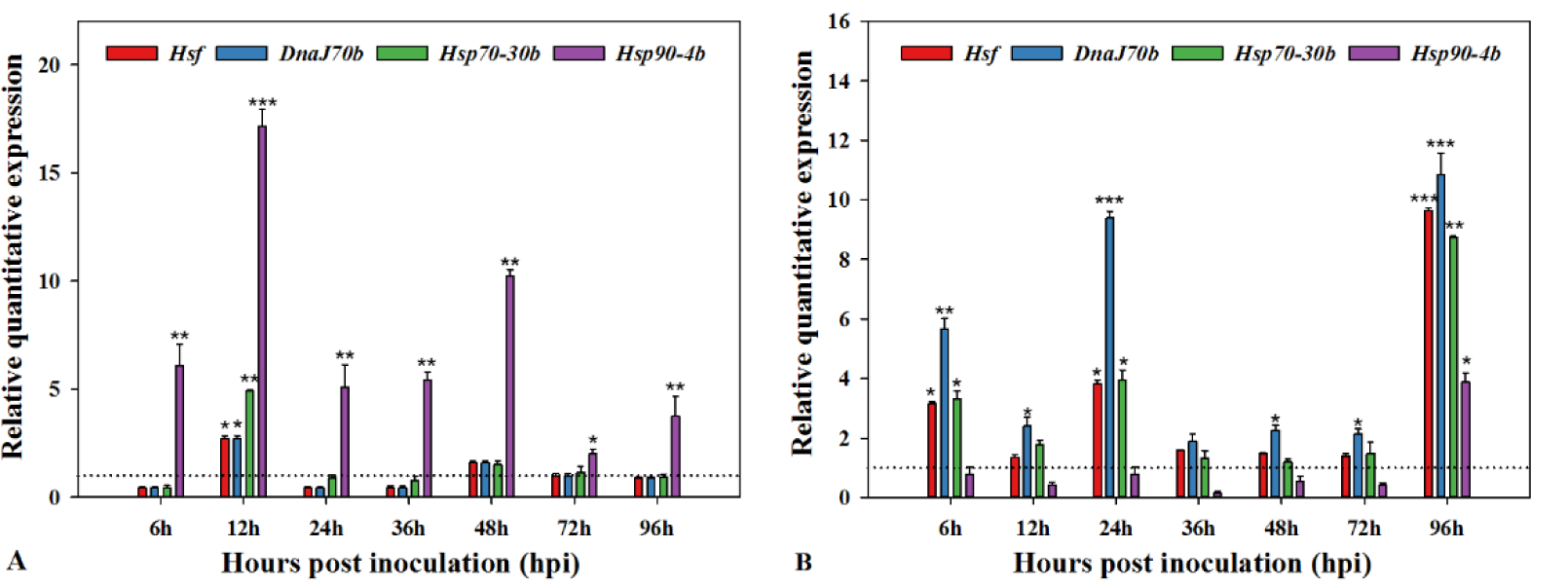
Co-expression patterns of selected heat shock transcription factor and HSP-encoding genes in N9134R and N9134S induction by powdery mildew pathogen. Gene expression levels were assessed by qRT-PCR at 6, 12, 24, 36, 48, 72, and 96 hpi. Data were normalized to the α-tublin expression level. The mean expression value was calculated from three independent replicates, and the standard deviation was given at each time points. A represents gene expression in N9134R with resistance to powdery mildew E09; B represents gene expression in susceptible line N9134S. The dotted line indicated the controls at the same time points. The names of corresponding gene were listed in the top of each panel.

### In vitro interactions among TaHsp70-30b, TaHsp90-4b, and TaDnaJ70b proteins

The *TaHsp90.2* and *TaHsp90.3* genes, located on the second and seventh partially homologous chromosomes, respectively, are involved in wheat responses to the stripe rust pathogen [21]. In *Arabidopsis thaliana*, The J-domain of DnaJ proteins reportedly interacts with Hsp70. Previously, TaDnaJ70, TaHsp70-30 and TaHsp90-4 were detected as specific induced genes in fungi stress [28]. To understand the roles of HSPs in wheat resistance, we tested the interaction between them using Y2H system. After the expression of the proteins of interest in the Y2H system was confirmed on synthetic dextrose (SD) medium lacking Trp and Leu, the yeast strains with BD- and AD-TaHsp were grown on SD/−Trp/−Leu/−His/−Ade medium. The TaDnaJ70 protein, either in full or with a truncated N-/C-terminal, failed to interact with TaHsp70-30 in the Y2H assay (**Figure 8A and 8B**). Similarly, TaHsp90-4 did not interact with TaDnaJ70 or TaHsp70-30 (**Figure 8A**). Thus, the interaction between DnaJ and Hsp70 may occur selectively or indirectly in wheat.

**Figure 8.**
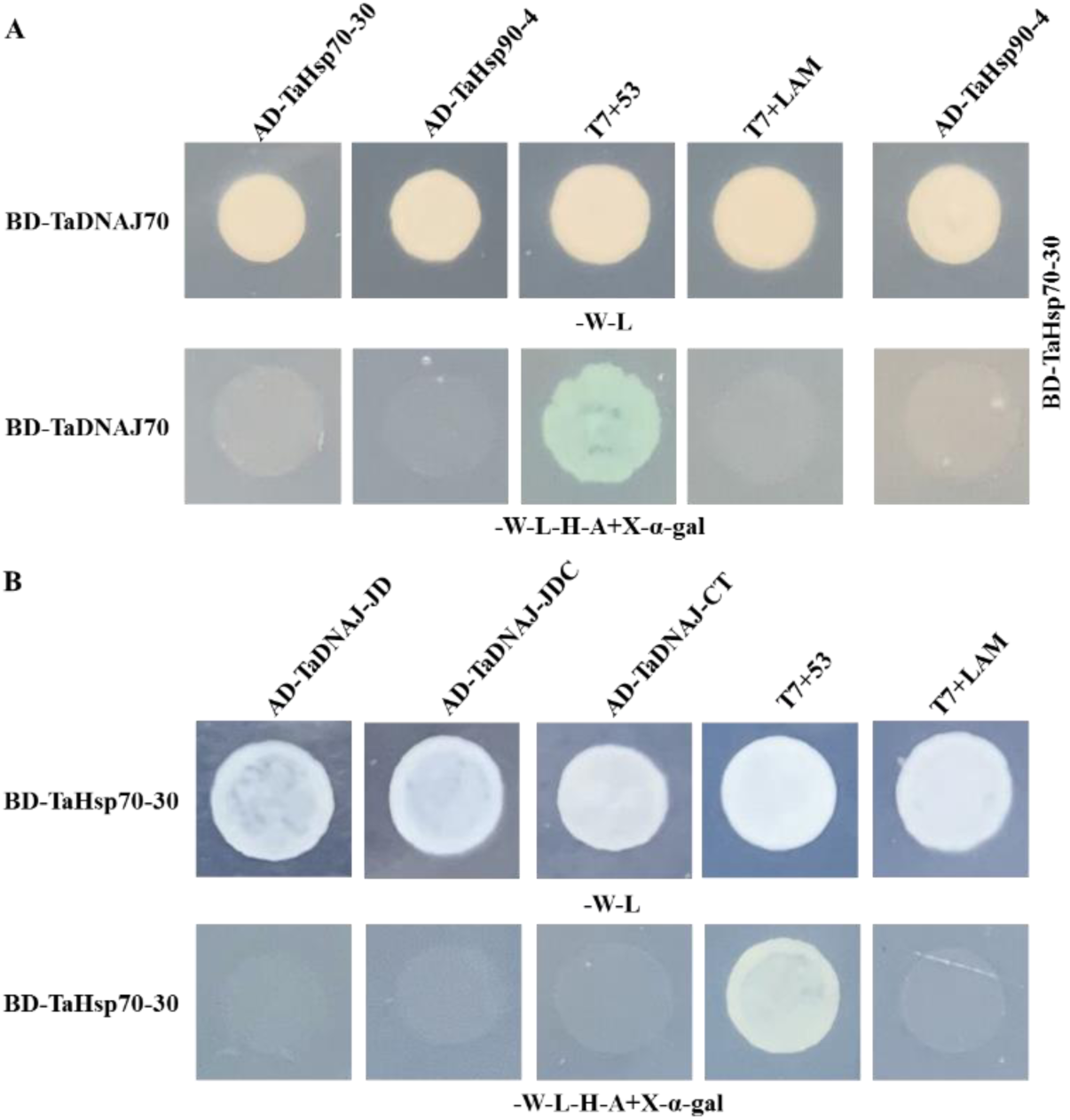
TaDnaJ70, TaHsp70-30 and TaHsp90-4 proteins interact with each other in a yeast two-hybrid system. A: TaDnaJ70 (a) or TaHsp70-30 (b) was cloned into pGBKT7 vector and the corresponding proteins were cloned into the pGADT7 vector. Cells of yeast strain AH109 harbouring the indicated plasmid combinations were grown on either the nonselective (SD-L-W) or selective (SD-L-W-H-A) medium containing 20 μg/mL X-a-gal. The interaction between SV40 large T-antigen (T) and murine p53 (53), T-AD+53-BD, was used as the positive control, while the interaction between T-antigen and human lamin C (Lam), T-AD+Lam-BD, was used as the negative control. A: interaction between full length proteins. B: interaction between full lengths TaHsp70-30 with the truncated DnaJ70 proteins. JD means the J and DRD domains; JDC represents the J, DRD and zinc finger domains, while CT means the C-termianl of TaDnaJ70.

## Discussion

### DNAJ protein groups, structure and classification

Historically, DnaJ proteins have been divided in three classes (I, II, and III) based on the G/F and zinc-finger motifs/domains present in *Escherichia coli* DnaJ proteins [33]. Obviously, the large structural and potential functional differences exist within and between Classes II and III. In this study, we detected 144 DnaJ family proteins, but only 9 out of 119 DnaJ have the classical CXXCXGXG domain. This domain was replaced by a CXXC repeat in a dozen DnaJ proteins, whereas the classical domain exists in 10 out of 13 DnaJ-CR proteins that lack the J-domain. Additionally, some of the proteins have similar N-terminal J-domain sequences, but are missing the aspartic acid in the HPD motif. For clarity, we refer to these domains as “J-like”. To clarify the biochemical functions or mechanisms associated with DnaJ proteins, we further classified these proteins in six groups with characteristic DRD, DED, CR zinc-finger or WAY motifs. The current study represents the first systematic analysis of wheat genes encoding Hsp40 and Hsp70 proteins, and generated useful information regarding the expression of HSP-encoding genes of common wheat. The presented data may form the basis for future investigations into the genome-wide *DnaJ* and *Hsp70* expression dynamics during plant–pathogen interactions. Furthermore, our findings may be relevant for further elucidating the role of HSPs in wheat defence responses as well as the detailed effects of the diverse HSP domains.

### Heat stress proteins play a critical role in wheat responding to pathogen

Heat shock proteins [e.g., Hsp40 (DnaJ), Hsp60, Hsp70, Hsp90, and Hsp101], which form one of the most ubiquitous classes of chaperones, have been implicated in diverse biological processes. The HSPs also directly stimulate cells of the innate immune system, suggesting they are activators of the innate immune system in animals [36, 37]. For example, a non-lethal heat shock induces Hsp70 synthesis and promotes the tolerance of shrimp to stresses due to heat, ammonia, and metals [38], and also prevents heart failure or ageing in humans [16]. So far, several plant species’ Hsp70s had been surveyed, such as *Hordeum vulgare* [25], *Brachypodium distachyon* [26], *A. thaliana* [39] and *Populus trichocarpa* [40]. Plant Hsp70s are localized to the cytosol, endoplasmic reticulum, mitochondria, chloroplasts, and peroxisomes [27]. Although there have been limited functional studies on plant Hsp70s, these proteins are believed to have multiple roles in plants and function like other eukaryotic Hsp70s. In *A. thaliana*, Hsp90 interacts with RAR1 and SGT1, which are major regulatory components of the disease resistance triggered by many R proteins [20, 41]. Similarly, Hsp90 proteins are required for the accumulation of potato Rx and tobacco N [42] as well as interactions with tomato I-2 [43] and NbSIPK [44], which regulate disease resistance. The available information related to the functions of HSPs in common wheat and other Triticeae species is still very limited. In wheat, *TaHSC70* encodes a TaHsp70-26 protein, and is responsive to stripe rust infections via a JA-dependent signal transduction pathway [45]. The cytosolic Hsp90s contribute to the resistance of wheat to the stripe rust fungus [21]. We previously identified Hsp70 as a key hub node protein, and determined that it likely plays a critical role in wheat defences against fungal infections [28]. In the current study, we identified 45 differentially expressed *DnaJ* genes as well as 14 *Hsp70* genes and six *Hsp90* genes in the fungus-inoculated N9134 seedling leaves. These results imply that HSPs influence plant resistance to fungi in ways that remain to be determined. Although it is unclear why the expression levels of HSP-encoding genes are considerably and differentially affected by powdery mildew and stripe rust infections in resistant and/or susceptible wheat leaves, the data presented herein verify that HSPs have diverse functions in plants.

### Heat stress proteins may indirectly interacted with each other

The Hsp90 chaperone pathway involves a series of steps, including the formation of multi-chaperone complexes with the assistance of receptors and cofactors. After Hsp40 binds to a receptor, its J-domain interacts with the C-terminus of Hsp70 [10]. The N-terminal of Hsp70 simultaneously binds to ATP. An intermediate complex, comprising Hsp70 and Hsp90, is then formed with the assistance of cofactors that help ATP bind to Hsp90 [46]. In this process, Hsp70 and Hsp90 are associated through the Hop adapter protein [12]. Additionally, because the chaperone partnership between Hsp40s and Hsp70s has been well established from yeast to humans [47], it was assumed, without experimental verification, that these HSPs also interact in plants. In fact, there are inter- and intra-species variations in the J-domain, hinting at the specificity of Hsp40–Hsp70 interactions [48]. In wheat, there are 144 DnaJ family proteins (119 DnaJ, 12 DnaJ-like, and 13 DnaJ-CR proteins) and 41 Hsp70s, with considerable diversity in both protein families. Thus, it is problematic if these proteins are able to freely interact with each other in wheat plants. In this study, our data indicated that TaDnaJ70 does not directly interact with either TaHsp70-30 or TaHsp90-4 or that our technique is not sensitive enough to detect the binding. There are two possible relationships between the DnaJ and Hsp70 proteins. Specifically, DnaJ may directly interact with specific Hsp70s, but not arbitrarily in wheat, or the interaction may be indirect (e.g., via an adapter), similar to the interaction between Hsp70 and Hsp90. Because the presence of the HPD motif in TaDnaJ70 has been confirmed, these results imply that other residues and regions outside the HPD motif contribute to the interaction between Hsp40 or Hsp40-like proteins and Hsp70. Our results provide the foundation for future wheat HSP studies for clarifying the interaction between HSPs and pathogen defence.

## Conclusions

Heat shock proteins (HSPs) play a crucial role in development and responses to diverse stresses. Here, we systematically classified DnaJ (Hsp40) proteins into six and four groups according to the detailed structural characterization, including HPD, DRD, DED, WAY, GK and CR domains. Moreover, infection by *Bgt* and *Pst* triggered robust alteration in gene expression of Hsp-encoding genes in *T. aestivum*, but the expression profiles of these HSP-encoding homologous genes varied in response to *Bgt* and *Pst*. The yeast two-hybrid assay experiments showed that a direct interaction are failed between TaDnaJ70, TaHsp70-30b and TaHsp90-4b.

These indicate that the Hsp protein-encoding genes of wheat responded to *Bgt* and *Pst* stress and played important roles in responding to fungal stress by a more complex pathway than that in mammal and model plant.

## Authors’ contributions

HZ and WJ conceived the project and provided overall supervision of the study; HG and GW performed the experiments and data analysis; YW and CW contribution to developing the materials; XL helped in experimental works; HZ wrote the first version of the paper; all authors reviewed and approved the final manuscript.

## Acknowledgments

This work was financially supported by the National Key Research and Development Program of China (grant no. 2017YFD0100701) and Academic Innovation and Intelligence Program in Higher Education Institutions. We thank Liwen Bianji, Edanz Editing China (www.liwenbianji.cn/ac) for editing the English text of a draft of this manuscript.

## Supporting information

Supplemental Table S1. Details regarding primers used for the qRT-PCR analysis and PCR amplification of HSP-encoding genes

Supplemental Table S2. Wheat DnaJ proteins

Supplemental Table S3. Wheat 70-kD heat shock proteins Supplemental Table S4. Wheat 90-kD heat shock proteins

Supplemental Figure S1. Expression patterns of differentially expressed *DnaJ* genes in N9134 infected by *Bgt* and *Pst*

Supplemental Figure S2. Expression patterns of differentially expressed *Hsp70* and *Hsp90* genes in N9134 infected by *Bgt* and *Pst*

Supplemental Figure S3. Relative expression of genes encoding *Bgt*-HSP during the wheat–*Bgt* interaction

