## Supplemental Table for "Identification and Expression Analysis of Heat Shock Proteins in Wheat Infected with Powdery Mildew and Stripe Rust"

**Table S1.** Details regarding primers used for the qRT-PCR analysis and PCR amplification of HSP-encoding genes.

| **Primer** | **Forward primer sequence** | **Reverse primer sequence** |
|---|---|---|
| qTaHsf-B1 | 5- CTTCACGGAATGGAGCAACG -3 | 5- GGCGGCTACTATTCTTCGTTTG -3 |
| qTaDNAJ70b | 5- TGAGTGACCCTGAGAAGCGTGAG -3 | 5- CACCACCTCCTCCACCCATT -3 |
| qTaHsp70-30b | 5- CAAGGGTGAGGACAAGCAGT -3 | 5- CCTCTGGGAATCATTGAAGTAA -3 |
| qTaHsp90-4b | 5- CAAGTCGGACCTCGTCAACA -3 | 5- GAAACCGACACCAAACTGCC -3 |
| BD-TaDNAJ70b | 5-CATGGAGGCCGAATTCATGTTCGGCCGCGCCC-3 | 5-CTGCAGGTCGACGGATTTACTGTTGTGCGCAC-3 |
| AD-TaHsp70-30b | 5-GGAGGCCAGTGAATTCATGGCGCCGACCAAGG-3 | 5-CTCGAGCTCGATGGATTTAGTCGACCTCCTCGA-3 |
| AD-TaHsp90-4b | 5-GGAGGCCAGTGAATTCATGGCGACGGAGACCGAGA-3 | 5-CTCGAGCTCGATGGATTTAGTCGACCTCCTCCA-3 |
| BD-TaHsp70-30b | 5-CATGGAGGCCGAATTCATGGCGCCGACCAAGG-3 | 5-CTGCAGGTCGACGGATTTAGTCGACCTCCTCGA-3 |
| AD-DNAJ70b-JD | 5-GGAGGCCAGTGAATTCATGTACTACGAGGTGCTCG-3 | 5-CTCGAGCTCGATGGATTTAATCATATATCTCACGC-3 |
| AD-DNAJ70b-JDC | 5-GGAGGCCAGTGAATTCATGTACTACGAGGTGCTCG-3 | 5-CTCGAGCTCGATGGATTTACTCCTGAAGAACCTTC-3 |
| AD-DNAJ70b-CT | 5-GGAGGCCAGTGAATTCATGCTTCAGGAGAAGAAAG-3 | 5-CTCGAGCTCGATGGATTTACAGCGAGTCGGGGAAC-3 |

**Table S2. Wheat DnaJ proteins**

| Number | Domains | Type | Gene ID in chromosome A | Gene ID in chromosome B | Gene ID in chromosome D | paralogue protein loci |
|---|---|---|---|---|---|---|
| **TaDNAJ 2** | J+DRD+CT | Ia | | | TraesCS1D02G004800 | |
| **TaDNAJ 3** | AR+DD+J | Vb | TraesCS1A02G020600 | TraesCS1B02G024600 | TraesCS1D02G020100 | |
| **TaDNAJ 4** | J+DRD+2C+WAY | Va | TraesCS1A02G042500 | TraesCS1B02G054900 | TraesCS1D02G042900 | |
| **TaDNAJ 5** | J+DRD+GF | IIb | TraesCS1A02G070300 | TraesCS1B02G088700 | TraesCS1D02G072800 | |
| **TaDNAJ 7** | J+DRD+GF+CT | IIb | TraesCS1A02G099300[P] | TraesCS1B02G125100[P,S] | TraesCS1D02G107200[P] | |
| **TaDNAJ 13** | J+DRD+CT | III | TraesCS1A02G175900 | TraesCS1B02G204300 | TraesCS1D02G162900 | |
| **TaDNAJ 9** | NT+J+LR+AS | Vb | TraesCS1A02G193300 | TraesCS1B02G208100 | TraesCS1D02G197000 | |
| **TaDNAJ 12** | J+DRD+AR+CT | III | TraesCS1A02G223900 | TraesCS1B02G237200 | TraesCS1D02G225500 | |
| **TaDNAJ 49** | NT+J+DRD+CT | IIa | TraesCS1A02G230000 | TraesCS1B02G244100 | TraesCS1D02G232000 | |
| **TaDNAJ 14** | GF+AP+J+DRD+CT | VI | TraesCS1A02G234900[P] | TraesCS1B02G248800[P] | TraesCS1D02G237300[P] | |
| **TaDNAJ 16** | AP+J+CT | III | TraesCS1A02G248900 | TraesCS1B02G259600 | TraesCS1D02G248400 | |
| **TaDNAJ 17** | AP+DD+J | VI | TraesCS1A02G251500 | TraesCS1B02G262100 | TraesCS1D02G251000 | |
| **TaDNAJ 18** | GF+J+DRD+GR+CT | IIb | TraesCS1A02G267600 | TraesCS1B02G278300 | TraesCS1D02G267500 | |
| **TaDNAJ 19** | AP+J+DRD+CT | III | TraesCS1A02G349900 | TraesCS1B02G364200 | TraesCS1D02G352600 | |
| **TaDNAJ 10** | J+DRD+G/F | Ib | TraesCS1A02G366400 | TraesCS1B02G384100 | TraesCS1D02G372000 | |
| **TaDNAJ 21** | J+DRD+GF+CT | IIb | TraesCS1A02G395300 | TraesCS1B02G423600 | TraesCS1D02G403500[P] | |
| **TaDNAJ 22** | AP+Un+J | VI | TraesCS1A02G419500 | TraesCS1B02G449600 | TraesCS1D02G427300 | |
| **TaDNAJ 23** | GR+J+DRD+CT | Vb | TraesCS1A02G420500[P] | TraesCS1B02G451500 | TraesCS1D02G428400[P] | |
| **TaDNAJ 24** | J+DRD+DD | III | TraesCS2A02G150400 | TraesCS2B02G175600 | TraesCS2D02G155500 | TraesCS6A02G059000LC |
| **TaDNAJ 25** | NT+J+DRD+CT | III | TraesCS2A02G160100 | TraesCS2B02G186400 | TraesCS2D02G167500 | |
| **TaDNAJ 26** | AP+J+DRD+CT | III | TraesCS2A02G272700 | TraesCS2B02G290900 | TraesCS2D02G272000 | |
| **TaDNAJ 27** | J+DRD+AP+2C+DD+CT | Va | TraesCS2A02G284200 | TraesCS2B02G301100 | TraesCS2D02G283100 | |
| **TaDNAJ 28** | SP+J+DRD+CT | Va | TraesCS2A02G498100 | TraesCS2B02G526300 | TraesCS2D02G498200 | |
| **TaDNAJ 29** | SP+J+DRD+AP+CT | III | TraesCS2A02G556100 | TraesCS2B02G586400 | TraesCS2D02G557300 | |
| **TaDNAJ 30** | NT+DD+J | VI | TraesCS2A02G576600 | TraesCS2B02G610800 | TraesCS2D02G587300 | |
| **TaDNAJ 31** | DD+J+DRD+CT | Ia | | TraesCS3B02G004700 | TraesCS3D02G008500 | TraesCSU02G039400[P] |
| **TaDNAJ 20** | AP+J+DRD+CT | Ia | TraesCS3A02G083700 | TraesCS3B02G098700 | | |
| **TaDNAJ 32** | J+DRD+GF+CT | IIb | TraesCS3A02G161200 | TraesCS3B02G192300 | TraesCS3D02G168500 | |
| **TaDNAJ 33** | 2GR+GF+J | VI | TraesCS3A02G187500[P] | TraesCS3B02G216900[P] | TraesCS3D02G191200[P] | |
| **TaDNAJ 34** | J+DRD+GS | III | | | TraesCS3D02G193500 | |

| | | | | | | |
|---|---|---|---|---|---|---|
| **TaDNAJ 72** | KH+GF+J+DRD+CT | III | TraesCS3A02G190100 | | TraesCS3D02G193600 | |
| **TaDNAJ 35** | AP+J+DRD+CT | IIa | TraesCS3A02G201300 | TraesCS3B02G225000 | TraesCS3D02G203800 | |
| **TaDNAJ 15** | J+DRD+CT | Ia | TraesCS3A02G202200 | | TraesCS3D02G201400 | |
| **TaDNAJ 36** | J+DRD+2C+DD+GF+DD+CT | Va | TraesCS3A02G213800[P] | TraesCS3B02G244000 | TraesCS3D02G215900[P] | |
| **TaDNAJ 37** | J+DRD+CT | Ia | TraesCS3A02G216800 | TraesCS3B02G247200 | TraesCS3D02G218800 | |
| **TaDNAJ 38** | J+DRD+CT | Ib | TraesCS3A02G254700 | TraesCS3B02G286700 | TraesCS3D02G255600 | |
| **TaDNAJ 39** | AR+J+DRD+CT | III | TraesCS3A02G282100 | TraesCS3B02G315800 | TraesCS3D02G282100 | |
| **TaDNAJ 40** | NT+J+CT | VI | TraesCS3A02G309700[S] | TraesCS3B02G152700 | TraesCS3D02G135400 | |
| **TaDNAJ 41** | J+DRD+GF+CT | IIb | TraesCS3A02G386400 | TraesCS3B02G418500 | TraesCS3D02G379400 | |
| **TaDNAJ 42** | J+DRD+2C+RR+DD+CT | Va | TraesCS3A02G435100 | TraesCS3B02G470100 | TraesCS3D02G518300LC | |
| **TaDNAJ 43** | J+DRD+CT | IIb | TraesCS3A02G520100 | TraesCS3B02G587800 | TraesCSU02G059600 | |
| **TaDNAJ 44** | J+DRD+2C+2DD+CT | Va | TraesCS3A02G536400[P] | TraesCS3B02G607500 | TraesCS3D02G541800 | |
| **TaDNAJ 45** | LK+J+DRD+GF | VI | TraesCS3A02G537600[P] | TraesCS3B02G603100[P] | TraesCS3D02G543100[P] | |
| **TaDNAJ 46** | GR+J+DRD+CT | Vb | TraesCS4A02G521400LC | | | |
| **TaDNAJ 47** | AR+J+AS+DD | Vb | TraesCS4A02G353900 | | | TraesCS5B02G518600 |
| **TaDNAJ 48** | NT+J | VI | TraesCS4A02G256200 | TraesCS4B02G058500 | TraesCS4D02G058400 | |
| **TaDNAJ 50** | NT+J+DRD+2C+DD+DD | Va | TraesCS4A02G214500 | TraesCS4B02G101500 | TraesCS4D02G095600LC | |
| **TaDNAJ 51** | AS+J+DRD+CT | VI | TraesCS4A02G203300 | TraesCS4B02G107200 | TraesCS4D02G104100 | |
| **TaDNAJ 52** | NT+J+DRD+CT | Ia | | TraesCS4B02G175400 | TraesCS4D02G177300 | |
| **TaDNAJ 53** | AS+J+DRD+CT | Ia | TraesCS4A02G116800 | TraesCS4B02G187600 | TraesCS4D02G189000 | |
| **TaDNAJ 54** | NT+J+DRD+GF+CT | Ia | TraesCS4A02G110600 | TraesCS4B02G193500 | TraesCS4D02G194500 | |
| **TaDNAJ 55** | NT+J+DRD+CT | III | TraesCS4A02G087200[P] | TraesCS4B02G217200[P] | TraesCS4D02G217500 | |
| **TaDNAJ 11** | SP+J+DRD+RR | Ia | TraesCS4A02G069100 | TraesCS4B02G226300 | TraesCS4D02G226900 | |
| **TaDNAJ 56** | SP+J+DRD+GF+4CC+CT | IIa | TraesCS4A02G064300 | TraesCS4B02G241700 | TraesCS4D02G241300 | |
| **TaDNAJ 57** | GS+J | VI | TraesCS4A02G041000 | TraesCS4B02G261900 | TraesCS4D02G261900 | |
| **TaDNAJ 58** | J+DRD+AP+2C+DD+J | Va | | TraesCS4B02G319100[P] | TraesCS4D02G315600[P] | TraesCS5A02G493100 |
| **TaDNAJ 59** | J+DRD+AR+CT | III | | TraesCS4B02G334200 | | |
| **TaDNAJ 60** | J+DRD+CT | Ia | | | TraesCS4D02G334100 | |
| **TaDNAJ 61** | AS+J+DRD+CT | IIa | TraesCS5A02G045300[P] | TraesCS5B02G049500 | TraesCS5D02G055400[P] | |
| **TaDNAJ 62** | J+DRD+CT | Ia | TraesCS5A02G061200[P,S] | TraesCS5B02G070900[P,S] | TraesCS5D02G072700[P,S] | |
| **TaDNAJ 63** | J+DRD+GF+CT | VI | TraesCS5A02G106000[P,S] | TraesCS5B02G114500[P,S] | TraesCS5D02G116700[P,S] | |
| **TaDNAJ 64** | SP+J+DRD+GF+4CC+CT | IIa | TraesCS5A02G115000 | TraesCS5B02G115900 | TraesCS5D02G125500 | |

| | | | | | | |
|---|---|---|---|---|---|---|
| **TaDNAJ 65** | SP+J+DRD+GF+4CC+CT | IIa | TraesCS5A02G117600 | | TraesCS5D02G129700 | |
| **TaDNAJ 66** | GR+J+DRD+CT | Ib | TraesCS5A02G190500 | TraesCS5B02G192700 | TraesCS5D02G200400 | |
| **TaDNAJ 67** | AS+J+DRD+CT | Ib | TraesCS5A02G190700 | TraesCS5B02G192500 | TraesCS5D02G200200 | |
| **TaDNAJ 68** | J+DRD+CT | Ia | TraesCS5A02G240700 | | | |
| **TaDNAJ 69** | J+DRD+AR+CT | Ia | TraesCS5A02G271700 | TraesCS5B02G272000 | TraesCS5D02G279400 | |
| **TaDNAJ 70** | J+DRD+GF+4CC+CT | IIa | TraesCS5A02G372900[P,S] | TraesCS5B02G374900 | TraesCS5D02G382400[P,S] | |
| **TaDNAJ 71** | DD+J | VI | TraesCS5A02G391600 | TraesCS5B02G396500 | TraesCS5D02G401400 | |
| **TaDNAJ 73** | AP+J+DRD+CT | Ia | TraesCS5A02G399900 | TraesCS5B02G404700 | TraesCS5D02G409800 | |
| **TaDNAJ 74** | J+DRD+GF+4CC+CT | IIa | TraesCS5A02G426100 | TraesCS5B02G428000 | TraesCS5D02G434100 | |
| **TaDNAJ 75-1** | AP+J+DRD+CT | III | | TraesCS5B02G496700 | | |
| **TaDNAJ 75-2** | AP+J+CT | III | | TraesCS5B02G496800 | 5DL:437178_AA1465520 | |
| **TaDNAJ 76** | AP+J+DRD+CT | III | TraesCS5A02G483600 | TraesCS5B02G496900 | TraesCS5D02G497200 | |
| **TaDNAJ 77** | GR+J | VI | TraesCS5A02G559800LC | TraesCS5B02G607100LC[P] | TraesCS5D02G420700[P] | |
| **TaDNAJ 78** | AP+J+DRD+CT | III | | TraesCS5B02G508500 | TraesCS5D02G508400 | TraesCS4A02G365000 |
| **TaDNAJ 79** | SP+J+DRD+CT | IIa | | TraesCS5B02G511600 | TraesCS5D02G512100 | TraesCS4A02G361300 |
| **TaDNAJ 80** | AP+J+GR+CT | Va | | TraesCS5B02G519100 | | |
| **TaDNAJ 81** | AR+J+AP+CT | Vb | | | TraesCS5D02G518400 | |
| **TaDNAJ 82** | J+DRD+SP+CT | IIb | TraesCS6A02G087000 | TraesCS6B02G115000 | | TraesCSU02G019500 |
| **TaDNAJ 83** | J+DRD+CT | III | | TraesCS6B02G114100 | TraesCS6D02G076700 | |
| **TaDNAJ 84** | J+DRD+CT | III | | 6BS:514431_AA1659570 | | U:645311_AA2143950 |
| **TaDNAJ 85** | J+DRD+CT | IIIb | TraesCS6A02G103600LC | | | |
| **TaDNAJ 6** | NT+J+DRD+CT | Ia | TraesCS6A02G158600 | TraesCS6B02G192700 | TraesCS6D02G153900 | |
| **TaDNAJ 86** | NT+J+SP+2C+DD+CT | Va | TraesCS6A02G167100[P] | TraesCS6B02G194500[P] | TraesCS6D02G155600[P] | |
| **TaDNAJ 87** | J+DRD+GF+4CC+CT | IIb | TraesCS6A02G250800 | TraesCS6B02G274600 | TraesCS6D02G232600 | |
| **TaDNAJ 88** | GR+J+DRD+CT | Ia | TraesCS6A02G264600 | TraesCS6B02G546300LC | TraesCS6D02G250300 | |
| **TaDNAJ 89** | AS+J+DRD+CT | Ia | TraesCS6A02G310200 | TraesCS6B02G340100 | TraesCS6D02G289600 | |
| **TaDNAJ 90** | NT+J+DRD+CT | Ia | TraesCS6A02G314500 | TraesCS6B02G344500 | TraesCS6D02G293800 | |
| **TaDNAJ 91** | SP+J+DRD+CT | Ia | TraesCS6A02G341900 | TraesCS6B02G373800 | TraesCS6D02G322200 | |
| **TaDNAJ 92** | AP+J+DRD+GF+4CC+CT | IIa | TraesCS6A02G372700 | TraesCS6B02G410600 | TraesCS6D02G356800 | |
| **TaDNAJ 93** | J+DRD+GR+CT | IIb | TraesCS6A02G421300 | | | |
| **TaDNAJ 94** | ASP+J+DRD+GF+4CC+CT | IIa | TraesCS7A02G051000 | | | |
| **TaDNAJ 95** | J+DRD+GF+CT | Ib | TraesCS7A02G159500 | TraesCS7B02G064000 | | |

| | | | | | |
|---|---|---|---|---|---|
| **TaDNAJ 96** | J+DRD+GF+CT | Ia | TraesCS7A02G159600 | TraesCS7B02G064100 | |
| **TaDNAJ 97** | J+DRD+GF+CT | Ia | | | TraesCS7D02G160400 |
| **TaDNAJ 1** | SP+J+DRD+GF+4CC+CT | IIa | TraesCS7A02G176600 | TraesCS7B02G081600 | TraesCS7D02G177900 |
| **TaDNAJ 98** | AP+J+DRD+CT | Ia | TraesCS7A02G194600 | TraesCS7B02G100200 | TraesCS7D02G196200 |
| **TaDNAJ 99** | NT+J+DRD+CT | VI | TraesCS7A02G213900 | TraesCS7B02G121200 | TraesCS7D02G216000 |
| **TaDNAJ 100** | AP+J+DRD+RR | Ia | TraesCS7A02G220300 | TraesCS7B02G127100 | TraesCS7D02G221800 |
| **TaDNAJ 101** | NT+J+DRD+GF+CT | III | TraesCS7A02G235100 | TraesCS7B02G133300 | TraesCS7D02G235100 |
| **TaDNAJ 102** | J+DRD+2G/F | Ia | TraesCS7A02G250300 | TraesCS7B02G140900 | TraesCS7D02G249000 |
| **TaDNAJ 103** | SP+J+DRD+RR | Ia | TraesCS7A02G271600 | TraesCS7B02G170400 | TraesCS7D02G271900 |
| **TaDNAJ 104** | J+DRD+GF+CT | IIb | TraesCS7A02G298000[P] | TraesCS7B02G188900 | TraesCS7D02G293900 |
| **TaDNAJ 105** | GR+J+DRD+CT | Ib | TraesCS7A02G305000 | TraesCS7B02G205400 | TraesCS7D02G300500 |
| **TaDNAJ 106** | SP+J+GF+DED | IV | TraesCS7A02G506000 | TraesCS7B02G694400LC | TraesCS7D02G494000 |
| **TaDNAJ 107** | GF+J+GF+DED | IV | TraesCS7A02G506100 | | TraesCS7D02G494100 |
| **TaDNAJ 8** | AS+J+DED+CT | IV | TraesCS7A02G506700 | | TraesCS7D02G494800 |
| **TaDNAJ 108** | SP+GR+J+DED | IV | | | TraesCS7D02G494900 |
| **TaDNAJ 109** | GS+J+DED | IV | | | TraesCS7D02G495200 |
| **TaDNAJ 110** | SP+J+GF+DED | IV | | | TraesCS7D02G495500 |
| **TaDNAJ 111** | GR+J+DED | IV | | TraesCS7B02G413400 | |
| **TaDNAJ 112** | NT+GF+J+DED+CT | IV | | TraesCS7B02G414800 | |
| **TaDNAJ 113** | AP+J+DED | IV | | TraesCS7B02G415000 | |
| **TaDNAJ 114-1** | GR+J+DRD+RR | Ia | | TraesCS7B02G445500 | TraesCS7D02G516600 |
| **TaDNAJ 114-2** | GR+J+DRD+RR | Ia | TraesCS7A02G528700 | TraesCS7B02G445600 | TraesCS7D02G516700 |
| **TaDNAJ 114-3** | GR+J+DRD+RR | Ia | TraesCS7A02G528800 | | TraesCS7D02G516800 |
| **TaDNAJ 115** | AP+J+DRD+RR | Ia | | 7BL:577619_AA1879700 | |
| **TaDNAJ 114-4** | GR+J+DRD+AR | Ia | TraesCS7A02G528900 | TraesCS7B02G445700 | TraesCS7D02G516900 |
| **TaDNAJ 114-5** | GR+J+DRD+AR | Ia | TraesCS7A02G529000 | TraesCS7B02G445800 | TraesCS7D02G517000 |
| **TaDNAJ 116** | NT+J+DRD+GR+CT | Ia | | TraesCS7B02G445900 | |
| **TaDNAJ 117-1** | GR+J+DRD+RR | Ia | TraesCS7A02G529400 | TraesCS7B02G446000 | |
| **TaDNAJ 117-2** | GR+J+DRD+RR | Ia | | TraesCS7B02G446100 | |
| **TaDNAJ 117-3** | GR+J+DRD+RR | Ia | | TraesCS7B02G448500 | |
| **TaDNAJ 118** | AP+J+DRD+CT | Ia | | TraesCS7B02G498100 | |
| **TaDNAJ 119** | SP+J+DED | IV | | TraesCSU02G098500 | |

| | | | | | |
|---|---|---|---|---|---|
| **TaDNAJ-CR 1** | GR+4CC | CR | TraesCS1A02G015100 | TraesCS1B02G019300 | 1DS:080446_AA0248140 |
| **TaDNAJ-CR 2** | AP+2CC | CR | TraesCS1A02G030000 | TraesCS1B02G035300LC | TraesCS1D02G030500 |
| **TaDNAJ-CR 3** | GR+4CC | CR | TraesCS2A02G058700 | TraesCS2B02G070700 | TraesCS2D02G057900 |
| **TaDNAJ-CR 4** | SP+4CC | CR | TraesCS2A02G136700 | TraesCS2B02G160600 | TraesCS2D02G139400 |
| **TaDNAJ-CR 5** | SP+4CC | CR | TraesCS2A02G136800 | | TraesCS2D02G139500 |
| **TaDNAJ-CR 6** | SP+PA+4CC | CR | TraesCS2A02G384200 | TraesCS2B02G401400 | TraesCS2D02G380900 |
| **TaDNAJ-CR 7** | AP+4CC | CR | TraesCS2A02G451100 | TraesCS2B02G472600 | TraesCS2D02G450800 |
| **TaDNAJ-CR 8** | GF+4CC | CR | TraesCS3A02G219600 | TraesCS3B02G250100 | TraesCS3D02G232900 |
| **TaDNAJ-CR 9** | RR+2CC | CR | TraesCS4A02G080900 | TraesCS4B02G223100 | TraesCS4D02G223500 |
| **TaDNAJ-CR 10** | GR+4CC | CR | TraesCS4A02G298500 | TraesCS4B02G015000 | TraesCS4D02G013200 |
| **TaDNAJ-CR 11** | SP+2CC | CR | TraesCS5A02G221700 | TraesCS5B02G220500 | TraesCS5D02G229500 |
| **TaDNAJ-CR 12** | AP+4CC | CR | TraesCS6A02G197700 | TraesCS6B02G218700 | TraesCS6D02G183000 |
| **TaDNAJ-CR 13** | AP+4CC | CR | TraesCS7A02G341000 | TraesCS7B02G242200 | TraesCS7D02G338600 |
| **TaDNAJL1** | JL+DRD+AR+CT | JL | TraesCS1A02G393800 | | |
| **TaDNAJL2-1** | GF+JL | JL | TraesCS1A02G419000[S] | TraesCS1B02G449200[S] | TraesCS1D02G426800[S] |
| **TaDNAJL2-2** | GF+JL | JL | TraesCS1A02G419100[S] | | |
| **TaDNAJL2-3** | GF+JL | JL | TraesCS1A02G419200[S] | | |
| **TaDNAJL3** | LA+JL+CT | JL | TraesCS2A02G095200 | TraesCS2B02G110300 | TraesCS2D02G093500 |
| **TaDNAJL4** | LA+JL+CT | JL | TraesCS2A02G155700 | TraesCS2B02G180900 | TraesCS2D02G161500 |
| **TaDNAJL5** | LA+JL+AP+1C+CT | JL | TraesCS2A02G165100 | TraesCS2B02G191300 | TraesCS2D02G172200 |
| **TaDNAJL6** | JL+DRD+CT | JL | TraesCS3A02G441600 | TraesCS3B02G475500 | TraesCS3D02G434400 |
| **TaDNAJL7** | LA+SS+JL | JL | TraesCS4A02G118900 | TraesCS4B02G185600 | TraesCS4D02G186900 |
| **TaDNAJL8** | GF+JL+DRD+1C+CT | JL | | TraesCS4B02G160100 | |
| **TaDNAJL9** | JL+DRD | JL | | TraesCS4B02G618600LC | |
| **TaDNAJL10** | AR+JL+DRD+CT | JL | TraesCS5A02G108400 | TraesCS5B02G114900 | TraesCS5D02G117100 |
| **TaDNAJL11** | JL | JL | TraesCS5A02G199900 | TraesCS5B02G198200 | TraesCS5D02G205500 |
| **TaDNAJL12** | AR+JL | JL | TraesCS5A02G245400 | | TraesCS5D02G252200 |

Note: Wheat DnaJ proteins were firstly named by the homologous genes in other plants, while the others were named according to the location in chromosome. The number of wheat DnaJ proteins was sorted by homologous group. The superscript letter ‘p’ and ‘s’ represent that the HSP-encoding gene was differentially expressed in powdery mildew and stripe rust respectively.

**Table S3. Wheat 70-kD heat shock proteins**

| Number | Gene ID in chromosome A | Gene ID in chromosome B | Gene ID in chromosome D | paralogue protein loci |
|---|---|---|---|---|
| **TaHsp70-1** | TraesCS1A02G120100[P] | TraesCS1B02G139500[P,S] | TraesCS1D02G121000[P,S] | |
| **TaHsp70-14** | TraesCS1A02G120200[S] | TraesCS1B02G139600 | TraesCS1D02G121200 | |
| **TaHsp70-6** | TraesCS1A02G133100 | TraesCS1B02G151300[S] | TraesCS1D02G131800 | |
| **TaHsp70-3** | | TraesCS1B02G163400 | | |
| **TaHsp70-4** | TraesCS1A02G285000[P,S] | TraesCS1B02G294300 | TraesCS1D02G284000[P,S] | |
| **TaHsp70-5** | TraesCS1A02G295600 | | | TraesCS2D02G575200 |
| **TaHsp70-7** | | TraesCS2B02G125400 | | |
| **TaHsp70-8** | TraesCS2A02G506900 | | | |
| **TaHsp70-9** | TraesCS2A02G507000 | TraesCS2B02G535000 | TraesCS2D02G625000LC | |
| **TaHsp70-10** | TraesCS2A02G585600 | | TraesCS2D02G600900 | TraesCS1D02G452500 |
| **TaHsp70-11** | TraesCS3A02G262900 | TraesCS3B02G296300 | TraesCS3D02G262800 | |
| **TaHsp70-12** | TraesCS3A02G305100 | TraesCS3B02G331400 | TraesCS3D02G296700 | |
| **TaHsp70-13** | | TraesCS3B02G390700 | TraesCS3D02G351900 | |
| **TaHsp70-15** | | TraesCS3B02G390800 | TraesCS3D02G352400 | |
| **TaHsp70-2** | TraesCS4A02G175100 | TraesCS4B02G142400[P,S] | TraesCS4D02G140800[P] | |
| **TaHsp70-18** | TraesCS4A02G098600 | TraesCS4B02G205700 | TraesCS4D02G206600 | |
| **TaHsp70-19** | TraesCS4A02G098200 | TraesCS4B02G206300 | TraesCS4D02G207100 | |
| **TaHsp70-20** | TraesCS4A02G098100 | | | |
| **TaHsp70-21** | | TraesCS4B02G206700 | TraesCS4D02G207500 | |
| **TaHsp70-22** | TraesCS4A02G066100 | TraesCS4B02G243400 | TraesCS4D02G243000 | |
| **TaHsp70-23** | | TraesCS4B02G397600 | | TraesCSU02G116400 |
| **TaHsp70-24** | | TraesCS4B02G545200LC | TraesCS4D02G338300 | TraesCS5A02G511700 |
| **TaHsp70-25*** | TraesCS5A02G078000 | TraesCS5B02G087700[P] | TraesCS5D02G093900 | TraesCS2B02G374700 |
| **TaHsp70-26** | TraesCS5A02G106200 | TraesCS5B02G111200 | TraesCS5D02G117600 | |
| **TaHsp70-27** | | TraesCS5B02G129700 | | |
| **TaHsp70-28** | TraesCS5A02G268100 | TraesCS5B02G267900 | TraesCS5D02G276100 | |
| **TaHsp70-29** | TraesCS5A02G298700 | TraesCS5B02G298000 | | |
| **TaHsp70-30** | TraesCS5A02G479300 | TraesCS5B02G492500 | TraesCS5D02G492900[S] | |
| **TaHsp70-31** | TraesCS6A02G042600 | TraesCS6B02G058300 | TraesCS6D02G049100 | |
| **TaHsp70-17** | TraesCS6A02G276700 | TraesCS6B02G304200 | TraesCS6D02G257000 | |

| | | | |
|---|---|---|---|
| **TaHsp70-25*** | TraesCS6A02G337100[P,S] | TraesCS6B02G367800[P,S] | TraesCS6D02G317700[P,S] |
| **TaHsp70-32** | TraesCS6A02G342200 | TraesCS6B02G374500 | |
| **TaHsp70-33** | TraesCS6A02G342400 | | |
| **TaHsp70-34** | TraesCS6A02G342500 | | |
| **TaHsp70-35** | TraesCS6A02G537000LC | | |
| **TaHsp70-36** | | | TraesCS6D02G322800 |
| **TaHsp70-37** | | | TraesCS6D02G339600 |
| **TaHsp70-38** | | | TraesCS6D02G339700 |
| **TaHsp70-39** | | | TraesCS6D02G339800 |
| **TaHsp70-40** | TraesCS7A02G430600 | TraesCS7B02G330900 | TraesCS7D02G422600 |
| **TaHsp70-16** | TraesCS7A02G457100 | TraesCS7B02G359100 | TraesCS7D02G445600 |
| **TaHsp70-41** | | TraesCS7B02G331400 | TraesCS7D02G423100 |

Note: Wheat Hsp70 proteins were named by the homologous genes in other plants at first, while the others were named according to the location in chromosome. The number of wheat Hsp70 proteins was sorted by homologous group. TaHsp70-25 was repeated and marked with star symbol because the protein could be encoded by both homologous group 5 and 6 in bread wheat. The superscript letter ‘p’ and ‘s’ represent that the HSP-encoding gene was differentially expressed in powdery mildew and stripe rust respectively.

**Table S4. Wheat 90-kD heat shock proteins**

| Number | Gene ID in chromosome A | Gene ID in chromosome B | Gene ID in chromosome D |
|---|---|---|---|
| **TaHsp90-1** | TraesCS2A02G033700 | TraesCS2B02G047400 | TraesCS2D02G033200 |
| **TaHsp90-2** | TraesCS5A02G101900[P] | TraesCS5B02G106300 | TraesCS5D02G113700[P] |
| **TaHsp90-3** | TraesCS5A02G251000 | TraesCS5B02G249000 | TraesCS5D02G258900 |
| **TaHsp90-4** | TraesCS5A02G260600 | TraesCS5B02G258900[P,S] | TraesCS5D02G268000 |
| **TaHsp90-5** | TraesCS7A02G242200[P,S] | TraesCS7B02G149200[P,S] | TraesCS7D02G241100[P,S] |
| **TaHsp90-6** | TraesCS7A02G529900 | TraesCS7B02G446900 | TraesCS7D02G517800 |

Note: The superscript letter ‘p’ and ‘s’ represent that the HSP-encoding gene was differentially expressed in powdery mildew and stripe rust respectively.

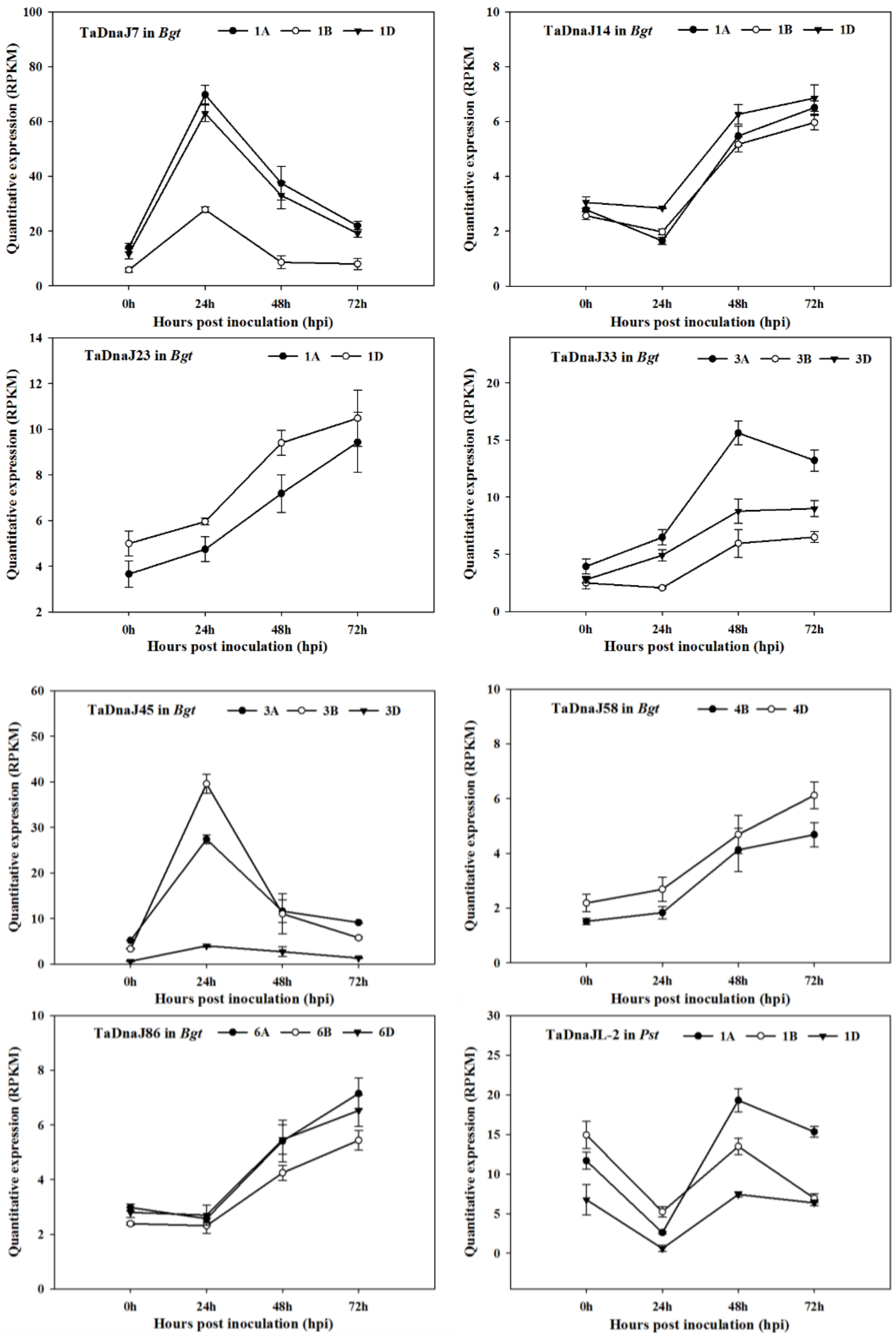


**Figure S1**. Expression patterns of differentially expressed *DnaJ* genes in N9134 infected by *Bgt* and *Pst*. Gene expression levels were assessed by transcript accumulation analysis. The mean expression value was calculated from three independent replicates. Line charts mean the gene expression of HSP-encoding gene in wheat infected by Pst and Bgt. The number 0, 24, 48 and 72 indicated the timepoints after infection. Bgt represents powdery mildew E09 inoculation condition; Pst represents stripe rust pathogen CYR 31 inoculation. The name of HSP-encoding genes and corresponding homologue genes were listed in the top of each panel.

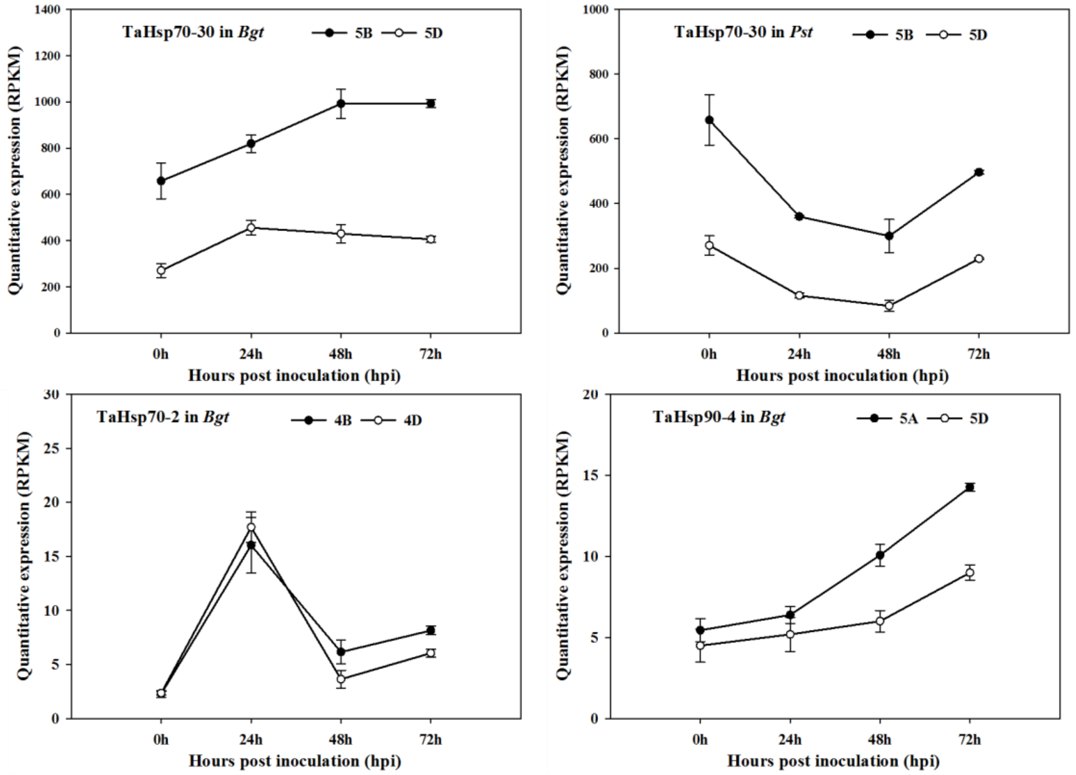


**Figure S2**. Expression patterns of differentially expressed *Hsp70* and *Hsp90* genes in N9134 infected by stripe rust and powdery mildew pathogen. Gene expression levels were assessed by transcript accumulation analysis. The mean expression value was calculated from three independent replicates. Line charts mean the gene expression of HSP-encoding gene in wheat infected by Pst and Bgt. The number 0, 24, 48 and 72 indicated the timepoints after infection. Bgt represents powdery mildew E09 inoculation condition; Pst represents stripe rust pathogen CYR 31 inoculation. The name of HSP-encoding genes and corresponding homologue genes were listed in the top of each panel.

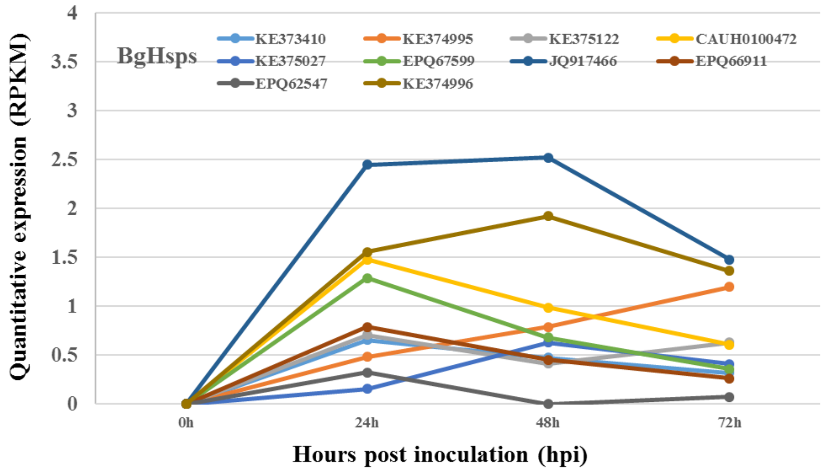


**Figure S3**. Relative expression of genes encoding Bgt-HSP during the wheat–*Bgt* interaction. Gene expression levels were assessed by transcript accumulation analysis. The mean expression value was calculated from three independent replicates. The X-axis indicated the timepoints at 0, 24, 48 and 72 hours after inoculation with powdery mildew Race E09. The names of homologue HSP-encoding genes of Bgt pathogen were listed in the top of panel with different color.
